# Protein-state dysregulation and sex-specific neurodevelopmental signatures in schizophrenia forebrain organoids

**DOI:** 10.64898/2026.06.01.729221

**Authors:** Helle Bogetofte, Sissel Ida Schmidt, Marie Sejberg Øhlenschlæger, Sofie Blomberg Elmkvist, Pia Jensen, Fadumo Abdullahi Mohamed, Anne With Mikkelsen, Elif Bayram, Jesper Foged Havelund, Matias Ryding, Lucrezia Criscuolo, Lene Andrup Jakobsen, Arkadiusz Nawrocki, Phillip Robinson, Madeline Lancaster, Jonathan Brewer, Nils Joakim Færgeman, Michael Eriksen Benros, Kristine Freude, Martin Røssel Larsen

## Abstract

Schizophrenia is highly heritable, yet the molecular mechanisms linking genetic risk to abnormal human brain development remain poorly understood. To address this, we generated dorsal forebrain organoids from 17 individuals with idiopathic schizophrenia and 17 age- and sex-matched controls and profiled them across multiple molecular layers, including single-nucleus transcriptomics, quantitative proteomics, metabolomics and deep post-translational modification (PTM) analysis. The organoids reproducibly modelled early cortical development and showed largely similar cellular composition between schizophrenia and control groups. Surprisingly, transcriptomic differences were relatively limited, with the strongest cell-type-specific changes observed in Cajal–Retzius neurons. In contrast, proteomic and particularly PTM-level analyses revealed widespread molecular disruption affecting pathways involved in neuronal migration, neurite development, synaptic function, protein kinase signalling, extracellular matrix organisation and lipid metabolism. Many of the earliest disease-associated changes emerged at the level of protein phosphorylation, consistent with altered neuronal maturation and neurite dynamics. At later developmental stages, schizophrenia organoids showed reduced abundance of synaptic proteins, fewer synaptic puncta and evidence of dysregulated retinoic acid and YAP1 signalling. Notably, most disease-associated alterations occurred independently of changes in transcript or protein abundance, indicating that key aspects of schizophrenia biology are encoded in protein state rather than expression level. These findings identify sex-specific dysregulation of protein state as a major molecular feature of schizophrenia and demonstrate the value of multi-layer proteomic approaches for uncovering disease mechanisms missed by transcriptomics alone.

## Introduction

Schizophrenia (SCZ) is a debilitating psychiatric disorder with a high heritability of 70–80%, characterized by a complex polygenic architecture^1,2^. Genome-wide association studies (GWAS) have identified hundreds of risk loci, the vast majority of which reside in non-coding regions and implicate broad biological pathways rather than single causal genes^2,3^. This polygenic distribution complicates mechanistic interpretation and limits the utility of conventional single-gene knockout models for capturing core disease biology. Instead, SCZ risk is thought to emerge from the cumulative effect of numerous modest genetic contributions, necessitating experimental systems that preserve the full patient-specific genomic context and polygenic burden^4^. Consequently, a fundamental challenge remains in understanding how polygenic risk disrupts specific cellular programs during human brain development - a period widely regarded as the critical window for SCZ pathogenesis^5^.

Human induced pluripotent stem cell (iPSC)-based models enable the study of patient-derived neural tissue while preserving endogenous genetic context^6^. Guided dorsal forebrain organoids (FOs) recapitulate key features of early human corticogenesis, including progenitor expansion, neuronal differentiation, and early circuit formation, providing experimental access to developmental stages that cannot be examined in postmortem tissue^7,8^. Molecular studies of SCZ iPSC models have, however, relied primarily on transcriptomic profiling^9–12^. Although informative, transcript abundance incompletely reflects cellular state, as mRNA levels often correlates poorly with protein abundance and do not capture post-translational regulation of protein activity and signaling dynamics^13^. These processes are central to brain development, where proliferation, migration, neurite extension, synaptogenesis, and metabolic remodeling depend on tightly regulated signaling networks and post-translational modifications (PTMs)^14^. Mass spectrometry-based proteomics and PTMomics therefore offer a more direct and functionally informative readout of phenotypes and disease biology not captured at transcript level. Yet, comprehensive proteomic interrogation of patient-derived brain organoids in SCZ has remained limited, particularly at scales sufficient to address inter-individual heterogeneity.

Here, we combined guided dorsal FOs derived from a large cohort of 17 individuals with SCZ and 17 age- and sex-matched controls (CTL) with an integrative multi-omics framework comprising single-nucleus transcriptomics, metabolomics, proteomics, deep PTM profiling, and imaging. To our knowledge, this represents one of the largest SCZ brain organoid cohorts analysed to date and among the most extensive proteomic/PTM phenotypic characterizations of any psychiatric disease model. By integrating protein abundance with phosphorylation, S-palmitoylation, cysteine redox modifications, and sialylated N-linked glycosylation, we capture regulatory layers not accessible through transcript-centric analyses alone^15^. This design enables robust detection of convergent molecular phenotypes across genetically heterogenous individuals and provides a high-resolution map of early developmental dysregulation in SCZ. Based on our data, we propose that SCZ is fundamentally a neurodevelopmental disorder of protein-state regulation.

### A deeply phenotyped SCZ iPSC-derived dorsal forebrain organoid cohort

To investigate the molecular consequences of polygenic SCZ risk during early human neurodevelopment, we established a deeply phenotyped patient-derived iPSC cohort comprising 17 individuals diagnosed with idiopathic SCZ and 17 age- and sex-matched healthy CTLs (Fig. 1a–b, Table S1-2). All participants underwent extensive clinical characterization prior to inclusion, including structured psychiatric interviews, standardized symptom assessments, cognitive testing, neurological examination, digital phenotyping, and linkage to nationwide health registries and electronic health records (Fig. 1a). This extensive clinical characterization ensured rigorous clinical validation of SCZ diagnoses and careful exclusion of psychiatric morbidity in controls.

**Figure 1.**
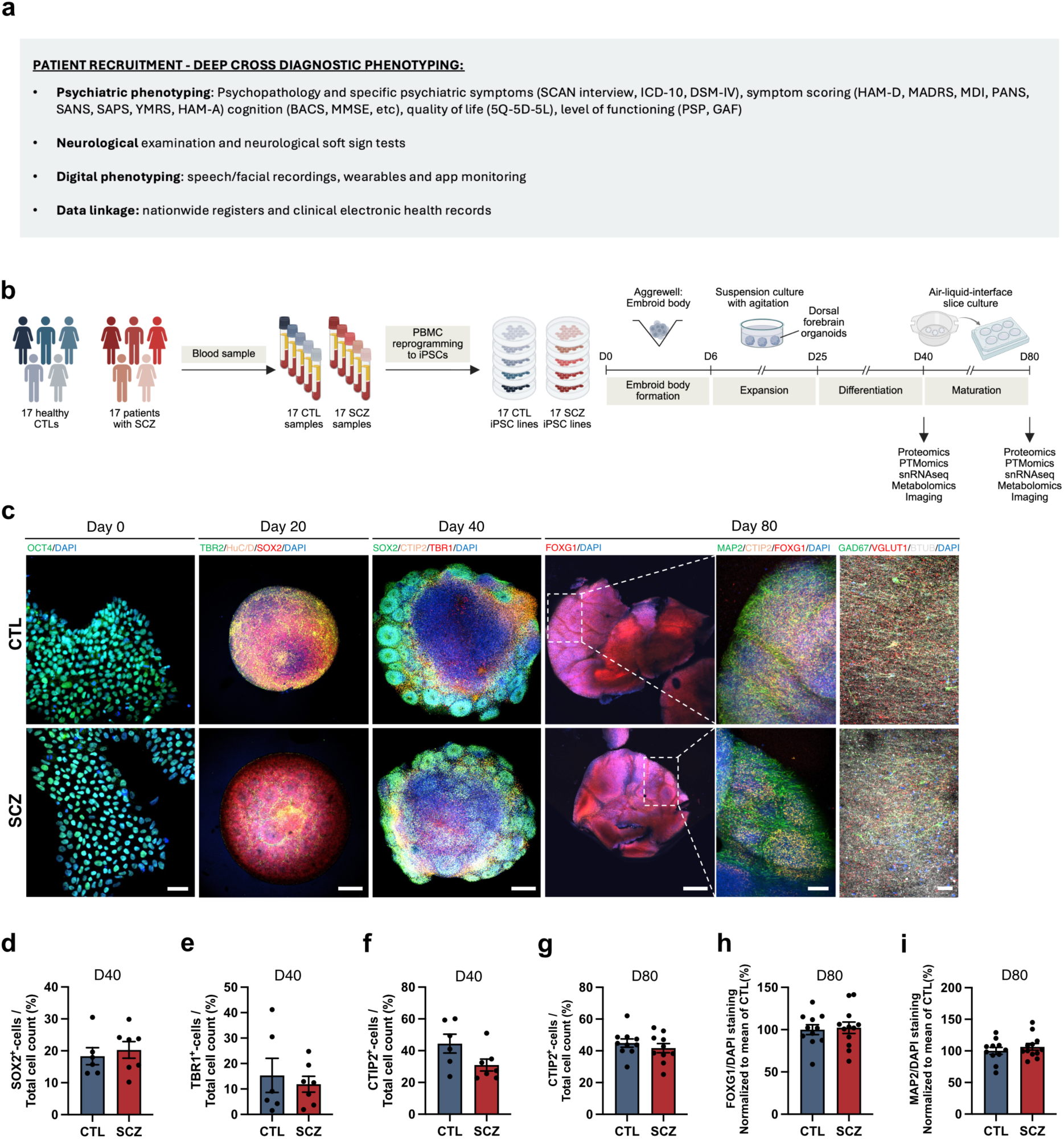
Establishment and characterization of a deeply phenotyped SCZ dorsal forebrain organoid cohort. **a.** Overview of the deep clinical phenotyping strategy used for cohort inclusion, including structured psychiatric assessment, cognitive testing, neurological examination, digital phenotyping, and linkage to nationwide health registries and electronic health records. Extensive clinical characterization was performed to ensure rigorous validation of schizophrenia (SCZ) diagnoses and exclusion of psychiatric disorders in healthy controls (CTL). **b.** Experimental workflow. Peripheral blood mononuclear cells (PBMCs) isolated from 17 individuals with SCZ and 17 age- and sex-matched healthy CTLs were reprogrammed into induced pluripotent stem cells (iPSCs), followed by dorsal forebrain organoid (FO) differentiation. Samples were collected at days 40 and 80 for multi-omic profiling and imaging analyses. **c.** Representative confocal images of CTL and SCZ lines during dorsal forebrain differentiation. Day 0 iPSC cultures stained for OCT4 (green) and DAPI (blue). Scalebar = 50 µm. Day 20 FOs stained for TBR2 (green), HuC/D (yellow), SOX2 (red), and DAPI (blue). Scalebar = 250 µm. Day 40 FOs stained for SOX2 (green), CTIP2 (yellow), TBR1 (red), and DAPI (blue). Scalebar = 250 µm. Day 80 air-liquid interphase FO cultures (ALIFOs) stained for FOXG1 (red), MAP2 (green), CTIP2 (yellow), and DAPI (blue). Scalebar = 1 mm on FOXG1 overview images, and 250 µm on zoomed-in images. D80 ALI-FOs stained for GAD67 (green), BTUB (grey), VGLUT1 (red), and DAPI (blue). Scalebar = 50 µm. **d-f.** Quantification of SOX2^+^-, TBR1^+^- and CTIP2^+^-cells of total cell count (DAPI) at day 40. n = 7 SCZ and n = 6 CTL, 3 FOs per replicate. **g-i.** Quantification of CTIP2^+^-cells of total cell count and FOXG1 and MAP2 staining intensity of total cell staining intensity (DAPI) at day 80. n = 11 CTL and 12 SCZ, 3 ALI-FOs per replicate. Data are presented as mean ± SEM, two-tailed unpaired t-test.

Peripheral blood mononuclear cells (PBMCs) were isolated from blood samples and reprogrammed into iPSCs using non-integrating approaches (Fig. 1b). All generated iPSC lines underwent rigorous quality control (QC) prior to downstream differentiation (Fig. S1). Only high-quality iPSC lines fulfilling predefined QC criteria were advanced for organoid generation, thereby reducing technical variability across the cohort.

To model early corticogenesis, we differentiated iPSC lines into dorsal FOs using a commercially available directed differentiation kit from STEMCELL Technologies (Fig. 1b). At day 40 of differentiation FOs were sliced into 300 µm air-liquid interphase FO cultures (ALIFOs) using a vibratome to maintain nutrient and oxygen access to the interior regions of the FO, enabling long-term culture^16^. Samples were collected both at days 40 and 80 for multi-omics profiling and imaging analyses.

Robustness and reproducibility of organoid differentiation across all lines were evaluated using immunofluorescence staining, demonstrating highly consistent developmental progression in both CTL and SCZ FOs and ALI-FOs across major corticogenic stages (Fig. 1c, Fig. S2-3). At day 0, iPSC cultures uniformly expressed the pluripotency marker OCT4. By day 20 of differentiation, FOs displayed prominent neuroepithelial rosette-like structures enriched for SOX2^+^-neural progenitors, TBR2^+^-intermediate progenitors, and HuC/D^+^-early neurons, consistent with active neurogenic zone formation. At day 40, organoids developed increasingly organized ventricular zone-like architectures containing SOX2^+^-radial glia together with CTIP2^+^- and TBR1^+^-deep-layer cortical neurons. Upon maturation to day 80, organoids expressed the forebrain marker FOXG1 alongside the neuronal maturation marker MAP2 and retained CTIP2^+^-cortical neuronal populations. In addition, mature organoids contained both GAD67^+^-inhibitory GABAergic neurons and VGLUT1^+^-excitatory glutamatergic neurons embedded within extensive βIII-tubulin (BTUB)^+^-neuronal networks, consistent with progressive cortical maturation (Fig. 1c).

Quantitative analyses demonstrated highly comparable differentiation efficiency between CTL and SCZ FOs (Fig. 1d–i). Organoid growth dynamics from day 0–40 were similar between groups, with no differences in organoid area (Fig. S4). Likewise, proportions of SOX2^+^- and TBR1^+^-populations were comparable between CTL and SCZ organoids. Although CTIP2^+^-deep-layer neurons showed a non-significant trend towards reduction in SCZ organoids at day 40, this difference was absent by day 80, potentially indicating a subtle delay in early neuronal maturation that normalizes during later development (Fig. 1f-g). Consistently, FOXG1 and MAP2 immunoreactivity at day 80 was similar between groups, supporting broadly preserved corticogenesis and neuronal maturation in idiopathic SCZ organoids.

### Limited transcriptomic disease alterations

To identify potential SCZ-related differences in cellular composition and transcriptomic profiles, we performed single-nuclei RNA sequencing (snRNAseq) on day 40 FOs and day 80 ALIFOs (12 SCZs vs. 12 CTL, minimum 2500 cells per line per timepoint) (Fig. S5a). Corresponding to early cortical development, both progenitors and radial glia as well as migrating and newly formed neurons, both excitatory and inhibitory, were identified. The proportions shifted with increasing percentages of both deep and upper layer neurons from day 40 to day 80 and no discernible difference between SCZ and CTL at either timepoint (Fig. 2a-b, Fig. S5b-d). Pseudo-bulk analysis revealed no differentially expressed genes (DEGs) at day 40 and only two transcripts, ARAP3 and ARL9, significantly upregulated at day 80 (Fig. 2c, Fig. S5e). Both transcripts are involved in Rho-GTPase-mediated regulation of neurogenesis, migration and neurite outgrowth^17,18^, yet have not previously been connected to SCZ pathogenesis. A recent analysis of FOs from patients with autism spectrum disorder (ASD), found significant transcriptomic changes in genetic, but not idiopathic ASD^19^, supporting the limited findings in our idiopathic SCZ cohort.

**Figure 2.**
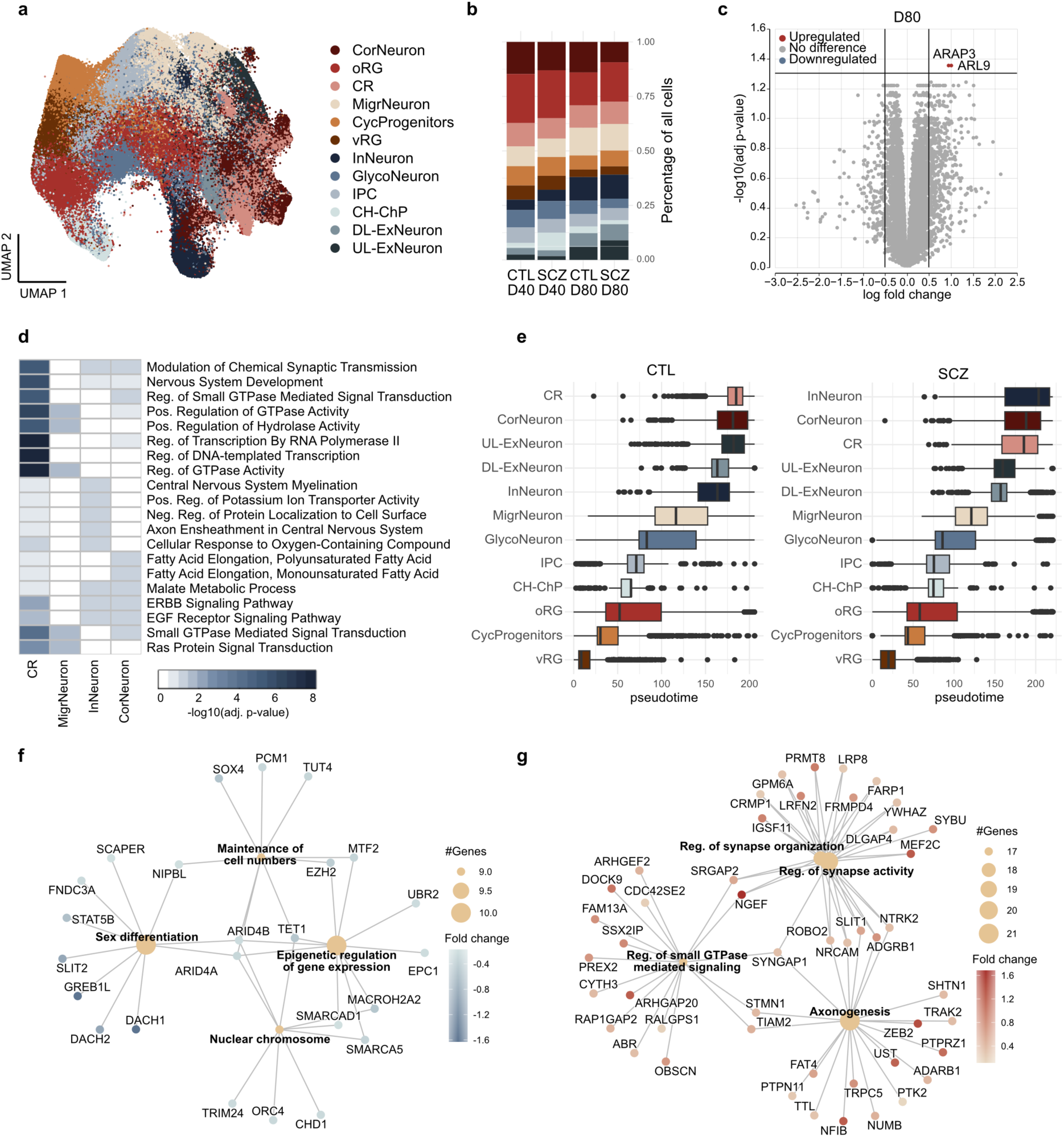
Limited transcriptomic SCZ changes, mainly affecting Cajal Retzius cells. **a.** UMAP embedding of day 40 and day 80 organoid single-nuclei RNA sequencing (snRNAseq) data with cells coloured by cluster/cell population: cortical neurons (CorNeuron), outer radial glia (oRG), Cajal Retzius cells (CR), migratory neurons (MigrNeuron), cycling progneitors (CycProgenitors), ventricular radial glia (vRG), inhibitory neurons (InNeuron), glycolytic neurons (GlycoNeuron), intermediate progenitors (IPC), cortical hem/choroid plexus (CH-ChP), deep layer excitatory neurons (DL-ExNeuron), upper layer excitatory neurons (UL-ExNeuron). **b.** Stacked barplot showing the proportion of each cell population for schizophrenia (SCZ) and control (CTL) organoids per time point. **c.** Volcanoplot of day 80 differentially expressed genes (DEGs) between SCZ/CTL. Limma voom-calculated adjusted p-value <0.05 and log-fold change >0.5 considered significant. **d.** Heatmap of top 20 enriched biological processes (BP) based on the four cell populations with significant DEGs. Fisher’s exact test with Benjamini-Hochberg corrected p-value. **e.** Pseudotime distribution by cell population for SCZ and CTL. **f-g.** Top four BPs in the CR population based on the significantly downregulated and upregulated DEGs. n = 11 SCZ and 12 CTL, four FOs or ALIFOs per replicate.

At day 40 transcriptomic profiles of the individual cell clusters were also identical. However, at day 80, significant cluster-specific DEGs could be identified in four of the neuronal clusters (Fig. 2d). The largest transcriptomic changes, with 365 DEGs, were identified in the Reelin+ Cajal Retzius (CR) population. Although pseudotime analysis overall pointed to similar developmental trajectories and timing comparing idiopathic SCZ and CTL, the CR population, together with the interneurons, showed slight differences in their developmental timing (Fig. 2e). This contrasts with results from 22q11.2 deletion SCZ patients cortical organoids that show abnormal pace of neuronal development^20^, again highlighting differences between genetic and idiopathic disease mechanisms. In the CR population, transcripts related to “nuclear chromosome” and “maintenance of cell numbers” were decreased whereas “regulation of synapse activity” and “axogenesis” were upregulated (Fig. 2f-g). These findings suggest the SCZ CR population to be at a more mature, less proliferative stage than CTLs. Although CR numbers are similar at this early stage, decreased levels could be a later consequence. This is notable given the importance of CR cells in establishing early cortical migration and reports of altered Reelin expression in SCZ postmortem^21–23^.

### Proteomic and PTMomic disease perturbations

To obtain a comprehensive characterisation of proteomic and metabolomic disease effects in SCZ organoids, we collected FOs and ALIFOs at day 40 and day 80 and extracted metabolites prior to performing our comprehensive PTMomic enrichment workflow (Fig.3a). We quantified 9605 and 8262 proteins at the two timepoints as well as peptides with phosphorylation (Phospho), N-linked sialylated glycosylation (Deglyco), lysine acetylation (LysAc) and free-(FreeCys), reversibly modified (RmCys) or S-palmitoylated cysteines (Depalm), respectively (Fig. 3a-b). This is to our knowledge the most exhaustive proteomic/PTMomic characterisation of patient-derived neural organoids to date. Around 40% of the identified proteins are listed as SCZ-associated in the Open Targets platform (Fig. 3b). A total of 88 and 100 differentially expressed proteins (DEPs) were significant between SCZ and CTL at the two timepoints, respectively, indicating protein perturbations to be more pronounced than transcript level changes. Around 64% of the DEPs were SCZ-associated, pointing to an enrichment of disease-related targets amongst these. However, a substantially larger number of proteins had significant PTM changes (Fig. 3c). Some proteins were affected both at the protein and PTM level, yet the majority of PTM level changes were found on proteins with similar abundance in SCZ and CTL (Fig. 3d). All PTM categories were represented amongst the significantly different with phosphorylation and cysteine-modifications constituting the largest proportions (Fig. 3e).

**Figure 3.**
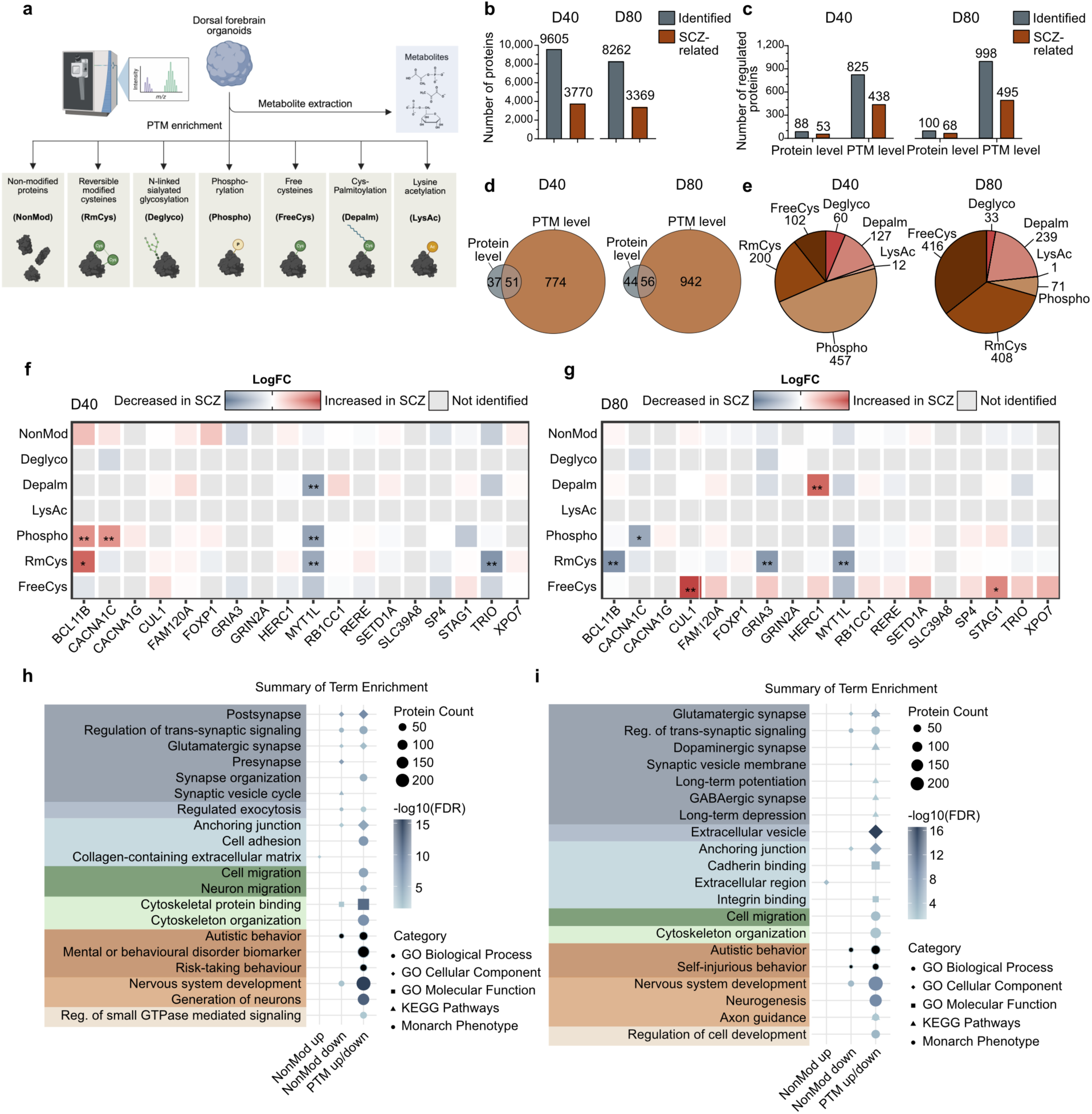
Proteomic and post-translational modification (PTM) changes in schizophrenia (SCZ) **a.** Experimental workflow for the proteomic, PTMomic and metabolomic analysis. **b.** Number of NonMod proteins identified per timepoint and how many of these were SCZ-associated. **c.** Number of significantly regulated proteins and proteins with significant PTM-level changes identified per timepoint and how many of these were SCZ-associated. Background-based t-test, n = 17, four FOs or ALIFOs per replicate, log2 fold change (SCZ/CTL) ≥ 0.263 and adjusted p-value ≤ 0.05 considered significant. **d.** Euler plot of overlap between proteins with significant NonMod abundance change and PTM-level change per timepoint. **e.** Proportion of each PTM type out of the total significantly changed PTMs. **f-g.** Average log2 fold change (SCZ/CTL) of protein abundance and PTM level peptide abundances for 18 protein-coding SCZ-associated GWAS-identified genes at day 40 and day 80. ** absolute log2 fold change (SCZ/CTL) ≥ 0.263, adjusted p-value ≤ 0.05, and abundances (grouped) CVs ≤ 60%, * absolute log2 fold change (SCZ/CTL) ≥ 0.263, adjusted p-value ≤ 0.05). **h-i.** Enrichment analysis on significantly regulated NonMod proteins (up-or down separately) and proteins with significant PTM regulation (up/down combined) showing the number of proteins connected to the term, the term category and the - log10(FDR) calculated by Fisher’s exact test with Benjamini-Hochberg correction for both timepoints.

Examining protein-coding SCZ-associated genes identified through GWAS-studies and rare variant analyses^2,24^, we found several with significant PTM changes at both timepoints (Fig. 3f-g). This included altered phosphorylation of BCL11b (S497), potentially altering its activation^25^ and altered levels of cysteine-modifications in CUL1 (C170), possibly affecting stability of the protein^26^ and in a zinc finger motif in MYT1L (C978), key to its function^27^. This highlights a potential role of common and rare genetic SCZ-associated variants in idiopathic forms of the disease through PTM dysfunction.

To understand which cellular pathways were affected in SCZ, we performed enrichment analyses on up- and downregulated proteins and PTMs (Fig. 3h-i). This emphasized that synaptic proteins, both glutamatergic, GABAergic and more general, are affected at both timepoints, especially with decreased protein abundance levels and PTM changes. Proteins related to cell migration, mental disorders and nervous system development showed similar changes. Extracellular matrix was the only enrichment identified based on proteins upregulated in SCZ. These findings align with pathway enrichment analyses results from a previous, more restricted proteomic study on SCZ patient and CTL cerebral organoids, pointing to decreased nervous system development proteins and an increased level of cell-cell adhesion proteins^28^. Overall, these results reveal converging disease pathways in idiopathic patient SCZ organoids through protein dysregulation.

### Early PTMomic changes converge on neuronal migration and synaptic maturation pathways

At day 40, DEPs in SCZ FOs prominently included glutamatergic receptor and synaptic proteins, including GRIA1, GRIA2, GRIA4 and GRM5, which were predominantly reduced relative to controls (Fig. 4a–b). Pathway enrichment analysis of DEPs identified chemical synaptic transmission as the top affected pathway, indicating early perturbations to synaptic development and neuronal communication (Fig. 4b). Across the integrated proteomic and PTMomic datasets, enriched pathways further included “neurexins and neuroligins”, “RHO GTPase cycle”, “L1CAM interactions”, and “extracellular matrix organization” (Fig. 4c). Collectively, these pathways regulate neuronal migration, neurite extension, axon guidance, and synaptogenesis during corticogenesis^29–31^, suggesting that early SCZ cortical development is characterized by coordinated disruption of structural and synaptic maturation programs.

**Figure 4.**
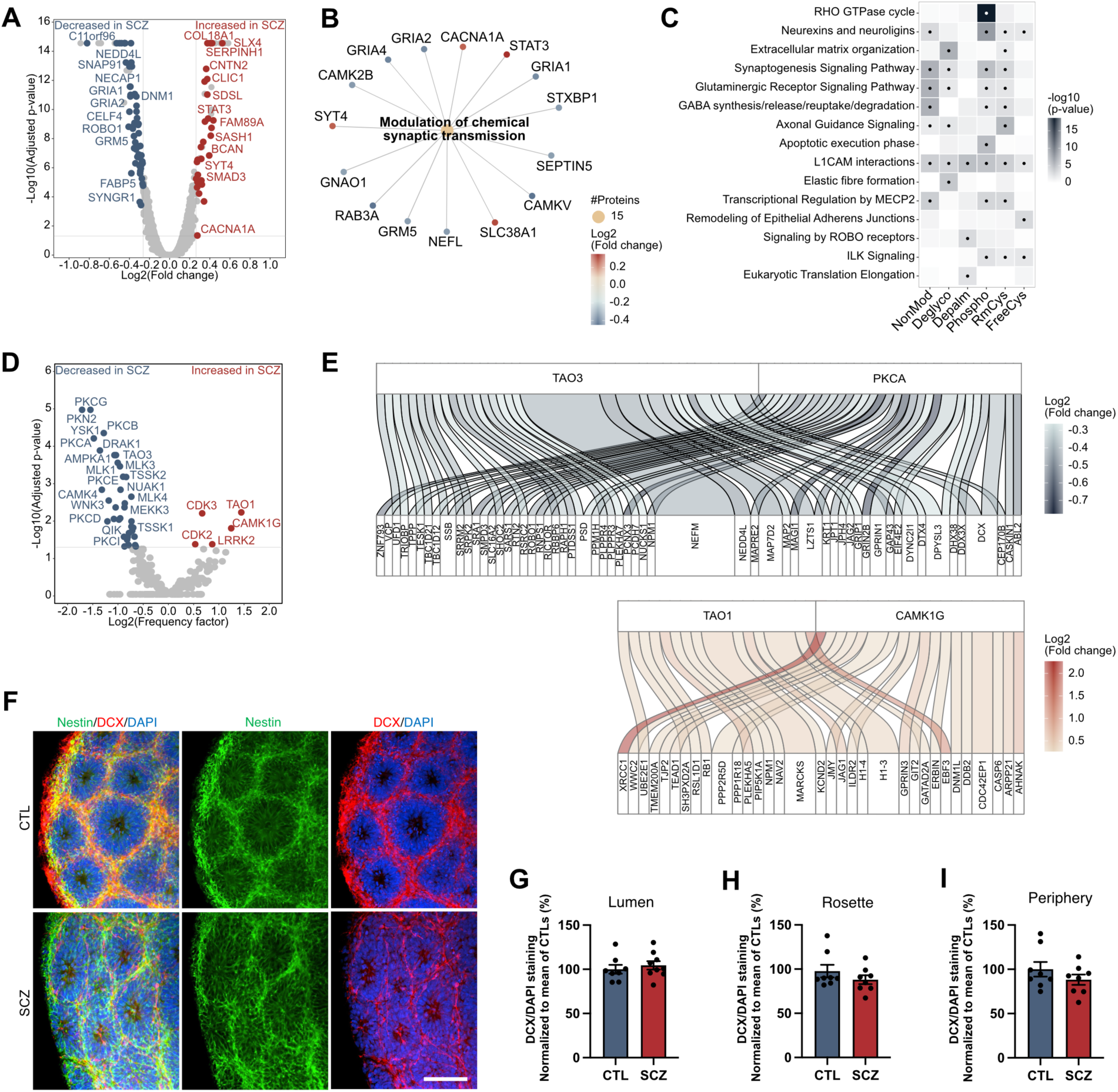
Early PTMomic changes to neuronal migration pathways. **a**. Volcano plot of day 40 differentially expressed proteins (DEPs) between schizophrenia (SCZ) and control (CTL) forebrain organoids. Background-based t-test, n = 17, log2 fold change (SCZ/CTL) ≥ 0.263, abundances (grouped) CVs ≤ 60% and adjusted p-value ≤ 0.05 considered significant. **b**. The top biological process enriched based on the significant DEPs with node colour indicating their log2 fold change. **c**. Heatmap of the top three biological pathways enriched based on significant DEPs or PTM peptides per dataset (non-modified proteins (NonMod), phosphorylation (Phospho), n-linked sialylated glycosylation (Deglyco), free-(FreeCys), reversibly modified (RmCys) or palmitoylated cysteines (Depalm)) with colour indicating term enrichment -log10(adjusted p-value) and dots adjusted p-value ≤ 0.05. **d**. Volcano plot of kinase activity prediction based on significant phospho-peptide changes in SCZ/CTL. One-sided Fisher’s exact test with Benjamini-Hochberg correction for multiple testing. Kinases with an absolute log2(Frequency factor) score ≥ 0.5 and adjusted p-value ≤ 0.05 considered significant. **e**. Key kinases with predicted decreased (top) and increased (bottom) activity in SCZ organoids and their protein substrates with significant phosphorylation changes. Flow size indicates the number of significant phospho-peptides per protein, and color indicates their average log2 fold change. **f)** Representative confocal images of CTL and SCZ day 20 FOs stained for Nestin (green), doublecortin (DCX, red), and DAPI (blue). Scalebar = 100 µm. g-i) Quantification of spatial distribution of DCX-positive immature neurons in and surrounding the neuroepithelial rosettes. n = 8 CTL and 9 SCZ, 3 FOs per replicate. Data are presented as mean ± SEM, two-tailed unpaired t-test.

With significant changes to phospho-peptides in many of the enriched pathways, we performed kinase activity prediction analysis based on differentially regulated phospho-peptides (Fig. 4d). This identified broad suppression of kinases associated with neuronal stabilization and synaptic maturation, including CAMK4, TAOK3, and multiple PKC family members^32–34^, whereas predicted activity of TAOK1, CAMK1G, CDK2, CDK3, and LRRK2 was increased in SCZ FOs. Several of these kinases converge on signaling pathways regulating neuronal polarization, microtubule dynamics, growth cone organization, and cortical neuron migration^33–36^, consistent with the enrichment of RHO GTPase and adhesion-related signaling pathways across the PTMomic datasets (Fig. 4c). Together, these findings suggest coordinated remodeling of opposing neuronal kinase states rather than isolated signaling defects. In particular, the reciprocal shift between increased TAOK1 and reduced TAOK3 signaling suggests dysregulation of a TAO kinase axis controlling neurite organization, microtubule stability, and cortical neuronal positioning^34,37^. Similarly, increased CAMK1G coupled with reduced CAMK4 activity is consistent with a shift in calcium-dependent signaling from neuronal stabilization toward cytoskeletal remodeling and neurite dynamics^38^. Notably, both TAO-family kinases, CAMK signaling components, and PKC-family members have previously been implicated in SCZ and related neurodevelopmental disorders, such as ASD^32,38–41^. Consistent with our findings, kinome profiling of DISC1-mutant SCZ iPSC-derived cortical neurons identified broad suppression of serine/threonine kinase activity, including altered AMPK, NUAK, WNK, CAMK1, and TAO kinase signaling^42^, supporting the hypothesis that dysregulated kinase networks represent a convergent feature of SCZ neurobiology.

Network analysis of kinase–substrate relationships further demonstrated coordinated phosphorylation changes in proteins regulating neuronal morphogenesis, cytoskeletal organization, and synaptic connectivity (Fig. 4e). Reduced predicted TAOK3 and CAMK4 activity converged on altered phosphorylation of proteins involved in radial migration, neurite stabilization, and glutamatergic synapse organization, including DCX, MAP2, NEFM, DPYSL3, SHTN1, ABI1, DLG2, GRIP1, and GRIN2B^33,34,37,38^. Conversely, increased predicted TAOK1 and CAMK1G activity preferentially affected phosphoregulation of proteins involved in neurite remodeling, polarity signaling, membrane dynamics, and early synaptic organization, including MARCKS, NAV2, GPRIN3, CDC42EP1, GIT2, and ERBIN^33,34,37,38^. Together, these findings support a model in which early SCZ cortical organoids exhibit a persistent neurite remodeling state coupled to impaired neuronal stabilization and synaptic maturation, further supported by broad suppression of PKC-family signaling linked to synaptic organization and activity-dependent neuronal stabilization.

Given that several affected kinase substrates, including DCX^43^, are central regulators of neuronal migration and neurite dynamics, we next examined the spatial distribution of DCX-positive immature neurons within day 20 FOs (Fig. 4f–i). Although overall DCX immunoreactivity was not significantly altered between CTL and SCZ organoids, SCZ organoids showed a subtle reduction in peripheral DCX-positive staining surrounding neuroepithelial rosettes. While not statistically significant, this trend is consistent with the predicted dysregulation of kinase networks involved in neuronal migration and neurite dynamics and may indicate mild delayed radial migration or maturation of newly generated cortical neurons.

### Validation of dysregulated synaptic proteins and upstream regulators

At day 80, a broader reduction of synaptic protein levels was identified (Fig. 5a, c) with trans-synaptic signaling being the top enriched term amongst downregulated DEPs. The top enriched term based on upregulated DEPs was “response to lipoprotein particle” based on MYD88, CD9 and APOE upregulation (Fig. 5b). This is notable given that disturbances in lipid metabolism are suggested contributors to SCZ pathogenesis through effects on membrane structure, neurotransmission, and neurodevelopment^44^. Across the proteomic and PTMomic datasets, “docasahexaenoic acid signaling” was also enriched (Fig. 5d). However, most terms were related to synapses with both glutamate- and GABAergic receptor signaling and overall synaptogenesis signaling affected (Fig. 5d).

**Figure 5.**
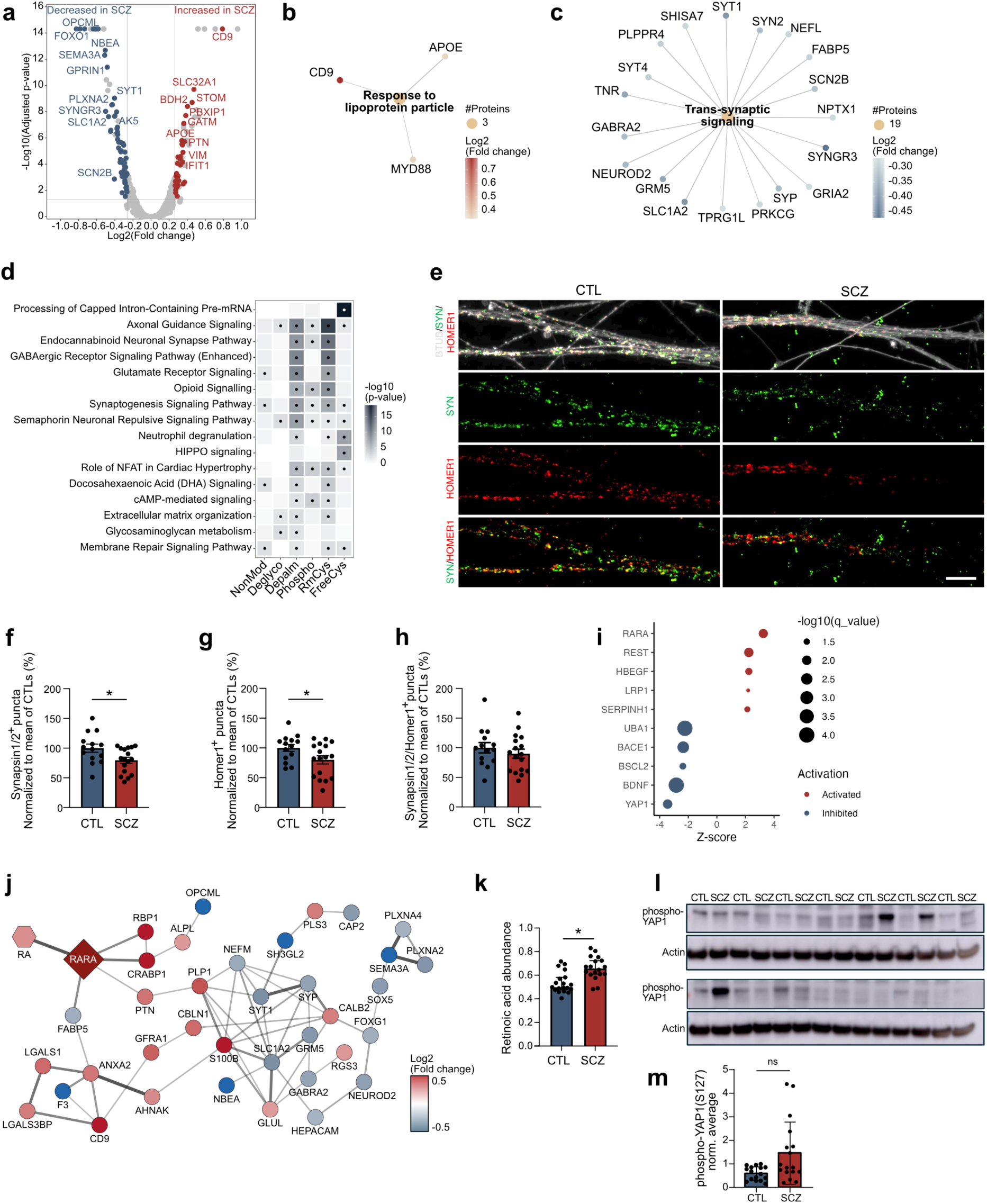
Proteomic/PTMomic changes to synaptic proteins result in lower synapse numbers with relevant predicted upstream regulators. **a.** Volcano plot of day 80 differentially expressed proteins (DEPs) between schizophrenia (SCZ) and control (CTL) air-liquid interface forebrain organoids (ALIFOs). Background-based t-test, n = 17, log2 fold change (SCZ/CTL) ≥ 0.263, abundances (grouped) CVs ≤ 60% and adjusted p-value ≤ 0.05 considered significant. **b-c.** The top biological processes enriched based on the significantly b. upregulated and c. downregulated DEPs with node colour indicating their log2 fold change. **d.** Heatmap of the top three biological pathways enriched based on significant DEPs or PTM peptides per dataset (non-modified proteins (NonMod), phosphorylation (Phospho), n-linked sialylated glycosylation (Deglyco), free-(FreeCys), reversibly modified (RmCys) or palmitoylated cysteines (Depalm)) with colour indicating term enrichment -log10(adjusted p-value) and dots adjusted p-value ≤ 0.05. **e.** Representative confocal images of CTL and SCZ day 80 ALI-FOs stained for synapsin 1/2 (SYN, green), Homer1 (red), BTUB (grey), and DAPI (blue). Scalebar = 10 µm. **f-h.** Quantification of synapsin 1/2^+^-puncta, homer 1^+^-puncta, and colocalized synapsin 1/2/homer1^+^-puncta relative to the mean of CTLs. n = 14 CTL and 17 SCZ, 3 ALI-FOs per replicate. Data are presented as mean ± SEM, two-tailed unpaired t-test, *adjusted p-value ≤ 0.05. **i.** Upstream regulators predicted based on significant DEPs with dot size indicating enrichment -log10(adjusted p-value) and colour predicted activation and inhibition. **j.** Network of significant DEPs (circles) connected to retinoic acid receptor alpha (RARA, square) and retinoic acid (RA, hexagon) with node colour indicating protein and metabolite log2 fold change. **k.** Retinoic acid abundance in CTL and SCZ ALIFOs measured by metabolomics. Student’s t-test with Benjamini Hochberg-based correction for multiple testing. n = 19 CTL and 18 SCZ, *adjusted p-value ≤ 0.05. **l-m.** Representative l. western blots and m. quantification of phospho-YAP1 (S127) in CTL and SCZ ALIFOs, normalized to beta-actin and average of all samples per blot. Mean ± SEM, n = 16 CTL and 17 SCZ, Mann-Whitney U-test.

To validate the observed synaptic protein dysregulation at the structural level, we performed immunocytochemical analysis of day 80 ALIFOs using the presynaptic marker synapsin 1/2 and the postsynaptic marker homer1 (Fig. 5e). Consistent with the proteomic findings, SCZ ALIFO neurites showed reduced density of synapsin 1/2^+^- and Homer1^+^-puncta relative to CTLs (Fig. 5f–g), whereas the number of colocalized synapsin 1/2/Homer1^+^-puncta was not significantly altered (Fig. 5h). Together, these findings support impaired synaptic protein abundance and synaptic maturation in SCZ organoids.

Based on the DEPs, we could predict upstream regulators likely responsible for the proteomic changes. Retinoic acid receptor alpha (RARA) was identified with a strong predicted activation z-score based on several of both up- and downregulated DEPs (Fig. 5i-j). Genetic and biochemical disruptions to retinoic acid (RA) signaling is well-established in SCZ and systemically increased RARA levels have been reported^45^. Supporting this, the metabolomic analysis identified RA levels as significantly increased in the SCZ ALIFOs (Fig. 5k). Disruptions in RA signalling have broad implications for neurodevelopment and could cause the synaptic connectivity changes identified^45^.

The upstream regulator with the largest inhibition z-score was the transcriptional co-activator YAP1 (Fig. 5i), which is required for proliferation and organization of radial glia during development^46^. Altered YAP1 transcription levels have recently been identified in postmortem SCZ brains^47^. YAP1 activity was evaluated using western blotting for YAP1 phosphorylated at S127, an inhibitory site. Although the average levels were not significantly different, several SCZs showed high levels of phospho-YAP1, indicating YAP1 dysregulation to be present in a subset of SCZ ALIFOs (Fig. 5l-m).

Combined, these results confirm the presence of synaptogenesis deficits in SCZ ALIFOs and disturbances to key upstream regulators, which are implicated in SCZ pathogenesis and could be causative.

### Sex-specific proteomic and PTMomic differences in SCZ ALIFOs

Examining the proteomic and PTMomic data on PCA plots, we observed a tendency for male and female samples to separate (Fig. 6a). We therefore reanalysed data separated into male and female. Surprisingly, the number of significant DEPs was substantially higher for both the female and male SCZ/CTL comparisons than when analysing all samples together. Furthermore, less than 15% of the DEPs were shared between females and males (Fig. 6b-d). Consistent with this, the upregulated DEPs resulted in different pathway enrichments between female and male SCZ/CTL. For female SCZ, “regulation of neurotransmitter levels” and “-metabolism” was affected based on increased levels of MAOA, MAOB, GAD1, COMT and ZNF219 amongst others. “Immune effector process” was also enriched based on several proteins including STAT3, PTX3 and CLU. For the male SCZ, proteins related to “double-strand break repair” mechanisms, which can be induced by oxidative stress, was enriched based on increased levels of MMC3, −4, −6 and −7. Additionally, “Tube/epithelium development” was enriched based on amongst others NFIB and NFIA, which are important transcription factors for forebrain development^48^. However, for both male and female SCZ, most DEPs were downregulated and enriched for similar GO terms for both sexes. The top enriched pathway for both male and female downregulated DEPs was “nervous system development”, yet very few DEPs were overlapping between the sexes (Fig. 6F). This indicates dissimilar proteomic changes between female and male SCZ with converging pathway effects.

**Figure 6.**
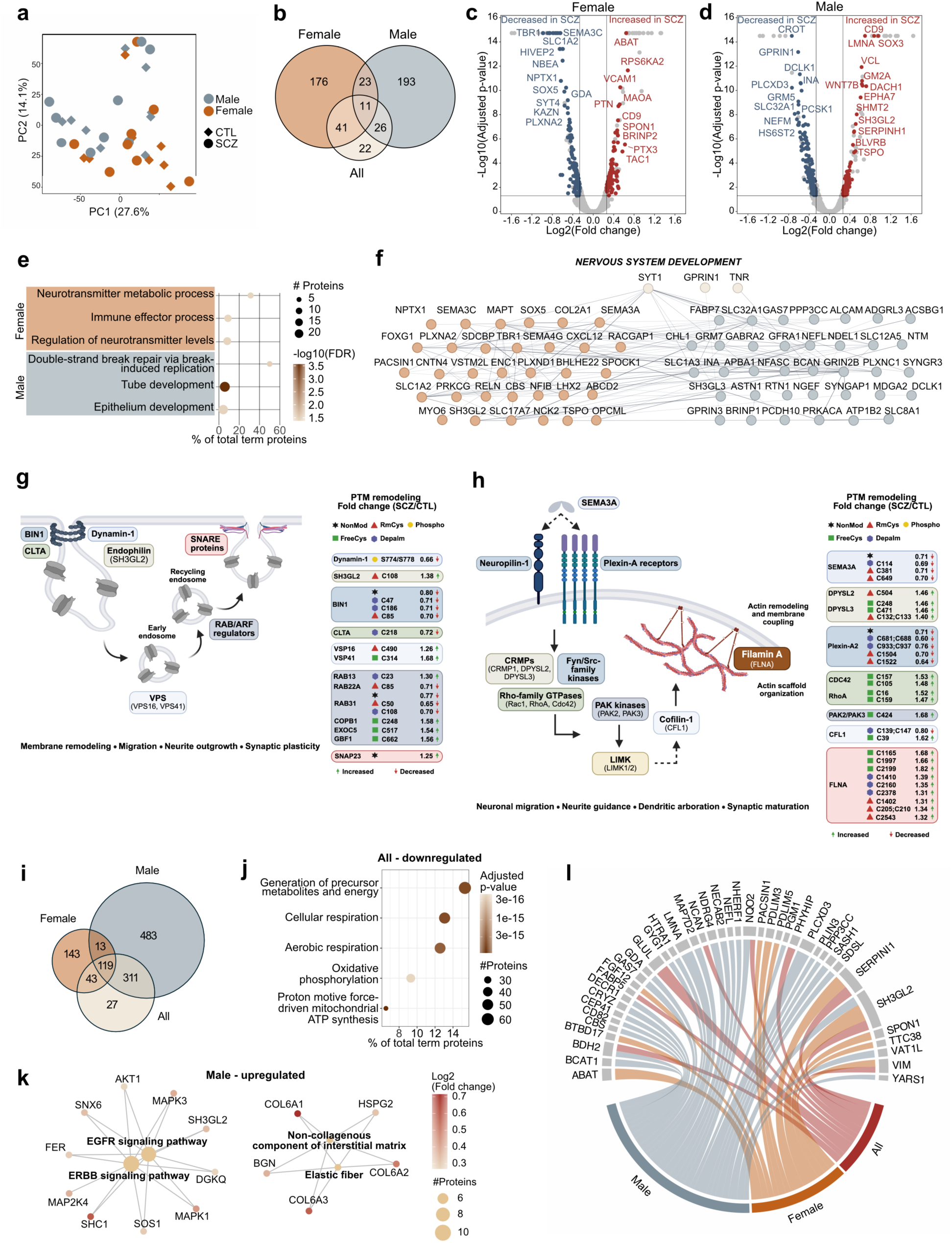
Sex-specific proteomic differences in schizophrenia (SCZ) forebrain organoids (FOs) and postmortem brains. **a**. Principal component plot based on nonmodified (NonMod) proteins from day 80 female (orange) and male (blue) SCZ (circles) and control (CTL, diamonds) air-liquid interface-FOs (ALIFOs). **b**. Euler plot of significant differentially expressed proteins (DEPs) comparing all SCZ to CTL ALIFOs (n = 17) or female (n = 8) or male (n = 9) alone. Background-based t-test, n = 17, log2 fold change (SCZ/CTL) ≥ 0.263, abundances (grouped) CVs ≤ 60% and adjusted p-value ≤ 0.05 considered significant. **c-d**. Volcano plot of day 80 DEPs between c. female SCZ and CTL and d. male SCZ and CTL ALIFOs. **e**. The top three GO terms enriched based on the significantly upregulated DEPs, analysing female and male separately, with node colour indicating enrichment -log10(adjusted p-value), size the number of term-related DEPs and the % of total term proteins on the x-axis. Fisher’s exact test with Benjamini-Hochberg correction. **f**. Significant DEPs between SCZ and CTL belonging to the term “nervous system development” identified in female (orange) or male (blue) or both (cream). **g-h.** Graphic illustration of g. neuronal dynamin-1 mediated endocytosis in male FOs and h. the SEMA3A signaling pathway in neurons in female FOs with log2(fold change) for NonMod proteins (black) or post-translational modifications (PTMs) marking the significant site and PTM-type: phosphorylatio (yellow), reversible cysteine-modifications (RmCys, red), free cysteines (FreeCys, green) and palmitoylation (Depalm, blue). **i**. Euler plot of significant DEPs in postmortem brains (Koopmans et. al 2026) comparing all (n = 47 SCZ, 29 CTL) or female (n = 24 SCZ, 22 CTL) or male (n = 23 SCZ, 27 CTL) alone. Unpaired t-test with Benjamini-Hochberg correction, n = 17, log2 fold change (SCZ/CTL) ≥ 0.263 and adjusted p-value ≤ 0.05 considered significant. **j**. The top five GO terms enriched based on the significantly downregulated DEPs in all postmortem SCZ vs. CTL cortex with node colour indicating enrichment -log10(adjusted p-value), size the number of term-related DEPs and the % of total term proteins on the x-axis. Fisher’s exact test with Benjamini-Hochberg correction. **k**. The top biological processes enriched based on the significantly upregulated DEPs in male alone with protein node colour indicating their log2 fold change and term node size indicating the number of connected DEPs. **l**. Chord diagram of DEPs identified in both SCZ vs. CTL ALIFOs and postmortem cortical tissue when analysis all, female or male samples alone. Size of chords indicate log2 fold change in ALIFOs.

Given these findings, we reanalysed the snRNAseq data comparing female and male SCZ and CTL separately (Fig. S6a-b). With a pseudobulk approach, no significant differences were present in females. This was not affected by X-inactivation erosion, which can occur in female iPSC lines^49^, as XIST was expressed in all lines and in comparable levels between SCZ and CTL (Fig. S6c). However, for the male samples a larger number of significant DEGs were identified both for the whole cell population and at the cluster level than when analysing all samples. The CR cells still displayed the largest number of DEGs, yet more cell populations were affected when analysing male SCZ and CTL alone, indicating that the transcriptomic changes are largely driven by the male patient ALIFOs (Fig. S6d).

At the PTMomic level we similarly identified sex-specific differences in which proteins and pathways were affected. In male SCZ FOs, a significant decrease in phosphorylation of dynamin-1 was identified on two important sites (S774 and S778), which when dephosphorylated allow binding to endophilin-A and initiation of endocytosis mechanisms. Several downstream proteins involved in neuronal endocytosis had significant PTM changes in male SCZ specifically (Fig. 6g). Altered endocytosis could have broad effects in SCZ neurons, affecting membrane remodelling, neuronal migration, neurite outgrowth and synaptic function^50^. In the female SCZ organoids, several proteins in the SEMA3A signaling pathway showed extensive significant PTM changes with potential effects on neuronal migration, neurite arborisation and dendrite formation (Fig. 6h)^51^.

### Sex-specific proteomic differences confirmed in postmortem SCZ patient prefrontal cortex

Although sex-specific transcriptomic/proteomic SCZ differences were indicated in a twin study of iPSC neurons, this has not previously been systematically demonstrated in a larger cohort of unrelated SCZ and CTL^52^. To investigate if sex-specific proteomic differences are also found in SCZ patient brains postmortem, we reanalysed recently published prefrontal cortex proteomics data from 47 SCZ patients and 49 CTLs^53^. Applying similar significance cutoffs as for our organoid data, we identified 500 significant DEPs in the cortical layer 4-6 data, which showed the highest degree of correlation with our ALIFO proteomics data (Fig. 6I, Fig. S7a-b). Comparing only female SCZ/CTL resulted in 318 DEPs whereas for males 926 DEPs were identified. This indicates substantial differences in the proteomic disease effects between female and male postmortem cortex, supporting the findings from SCZ patient ALIFOs. The significant downregulated DEPs for both female, male and all samples, strongly enriched for cellular respiration and mitochondrial energy production as originally reported (Fig. 6J, Fig. S7c-d)^53^. However, the upregulated DEPs, similarly to the ALIFO comparisons, resulted in different GO term enrichments: For all SZC samples, secretory vesicle lumen and extracellular matrix proteins were enriched in the upregulated DEPs (Fig. S7e). None were seen for the females, whereas for the males ERBB and EGFR signalling was identified based on AKT1, MAPK1, SH3GL2 and other DEPs (Fig. 6k). These findings are noteworthy in connection to the altered PTM-regulation of Dynamin-1 and endocytosis pathways in male SCZ ALIFOs. Dynamin-1 and endocytic regulation may precede changes in ERBB/EGFR signalling, which require receptor internalization for appropriate downstream activation^54,55^. Early abnormalities in Dynamin-1-dependent membrane trafficking could alter receptor availability, leading to secondary perturbations of ERBB/EGFR pathways.

When examining the intersection of DEPs in ALIFOs and postmortem cortex of all, female or male subjects, several notable proteins were identified (Fig. 6L). For all SCZ samples, BDH2 and SERPINI1^56,57^, which are both involved in oxidative stress defense, were significantly altered together with GLUL (Glutamine synthase), an important enzyme in glutamate homeostasis, which is decreased in SCZ cortex^58^.

However, the sex-specific comparisons resulted in larger numbers of common DEPs, highlighting the benefit of this approach for identifying disease-relevant perturbations. These proteins, showing differential expression in both early SCZ (ALIFOs) and late-stage (postmortem) disease stages, may be central to disease development. The one DEP, SH3GL2 (endophilin-A1), which was shared across all comparisons, is central to endocytosis-mediated synaptic vesicle recycling, dendritic spine morphogenesis, and neurotransmission^59,60^.

In conclusion, this study demonstrates that idiopathic SCZ can be modelled using patient-derived FOs and that proteomic/PTMomic approaches can identify disease perturbations with corresponding changes in postmortem patient cortical samples. Importantly, these results highlight extensive sex-specific proteomic/PTMomic disease profiles with both convergent and unique disease pathways affected, emphasising the need for sex-differentiated research and treatments into this disease.

## Materials and methods

### Recruitment of patients with Schizophrenia and age-matched healthy controls

The inclusion of patients with Schizophrenia (SCZ) and age-matched healthy controls (CTL) for generation of human induced pluripotent stem cell (hiPSC) lines is approved by The Regional Committee on Health Research Ethics in the Southern Region of Denmark (S-20220037) and by The Danish Data Protection Agency (P-2022-765). Participants came from the same cohort as described in^1,2^. Patients diagnosed with SCZ were recruited from psychiatric in- and out-patient clinics in the Capital Region of Denmark. Healthy CTLs were among others recruited via a Danish website for recruitment of participants to contribute to research studies (Forsøgsperson.dk). All participants had to be 18-50 years of age and came from the same community. All patients and healthy CTLs had thorough psychiatric evaluations including WHO Schedules for Clinical Assessment in Neuropsychiatry (SCAN, version 2.1), covering the last 4 weeks and performed by certified interviewers. For patients, the interview was used to validate the SCZ diagnosis given by clinical doctors prior to inclusion (according to the International Classification of Diseases, Tenth Revision (ICD10) F20), while for healthy CTLs the purpose was to rule out psychiatric disorders. Healthy CTLs were age- and gender-matched to the included patients with SCZ. hiPSCs were generated from peripheral blood mononuclear cells (PBMCs) derived from 34 individuals 19-29 years of age, including 17 diagnosed with SCZ and 17 healthy CTLs (Table S1-2).

Venous blood samples (≥34 mL) were collected from all participants into multiple 6 mL K2EDTA tubes (Greiner Bio-One GmbH) at the research facilities of the Copenhagen Research Centre for Biological and Precision Psychiatry, Mental Health Centre Copenhagen, Copenhagen University Hospital, and were immediately transported to the Department of Veterinary and Animal Science, University of Copenhagen, for further processing.

### Isolation of peripheral blood mononuclear cells and episomal reprogramming

PBMC isolation and reprogramming were performed as described in^1,2^. In brief, 34 mL of whole blood was diluted 1:1 with PBS−/− supplemented with 2% FBS and layered onto SepMate™-50 tubes containing Lymphoprep™ (#85450, #18060, Stem Cell Technologies (SCT)). Samples were centrifuged at 1200 xg for 10 min at room temperature (RT), after which the PBMC-containing top layer was collected, washed twice with PBS + 2% FBS, and centrifuged to remove residual contaminants.

Isolated PBMCs were cultured in cytokine-supplemented StemPro-34 medium (#10640,#10641, Gibco) prior to episomal reprogramming into hiPSCs using nucleofection (Lonza). Reprogrammed cells were cultured on Matrigel with sequential media changes until colony emergence around day 16. Expanded hiPSC clones were validated for pluripotency marker expression (Nanog, OCT-4, and TRA-1-60) by immunocytochemistry^2^.

The differentiation potential of hiPSC clones was assessed by embryoid body formation and spontaneous differentiation into the three germ layers, confirmed by immunocytochemistry for nestin (ectoderm), alpha-fetoprotein (endoderm), and smooth muscle actin (mesoderm)^2^. Genomic integrity was evaluated by copy number variation analysis using the CytoScan 750K array following DNA extraction from hiPSC clones (Fig. S1).

### Guided Dorsal Forebrain Organoid (FO) generation and air-liquid interphase FO cultures (ALIFOs)

hiPSCs were maintained in mTeSR Plus medium (#100-0276, SCT) on tissue culture-treated plates coated with Growth factor reduced Matrigel (#356230, Corning). Organoid generation was initiated when hiPSC cultures reached 70–90% confluency, displayed uniform morphology, and showed no signs of spontaneous differentiation. For passaging, cells were washed with calcium- and magnesium-free PBS (PBS −/−, #14190-094, Gibco) followed by incubation with Gentle cell dissociation (GCD) reagent (#100-0485, SCT) for 6-7 min at RT. The GCD solution was removed and the cell dissociated in fresh mTeSR Plus medium. Medium was changed every day. For thawing of cells, mTeSR Plus medium was added 10 µM ROCK inhibitor (#72304, SCT) for the first day.

Dorsal forebrain differentiation was initiated using the Dorsal Forebrain Organoid Differentiation Kit (#08620, SCT) according to the manufacturer until day 40 with minor modifications as described in the following. We used 50 µM ROCK inhibitor in the Seeding medium and seeded 2.5×10^6^ cells per Aggrewell^TM^ 800 (#34811, SCT). On day 6, cell aggregates were moved from the Aggrewells into a 15 mL Falcon tube using P1000 wide-bore pipette tips (#2079G, Thermo Scientific) and let to sink to the bottom. Medium was carefully removed and cell aggregates resuspended in 2 mL Expansion medium before being transferred to a low-attachment 60 mm dish (#430589, Corning). Cell aggregates were inspected under an EVOS^TM^ XL core imaging system (Invitrogen) in the sterile bench and 20-25 aggregates were transferred using a wide-bore P200 tip (#2069G, Thermo Scientific) to a well of a Nunc^TM^ non-treated 6-well plate (#150239, Thermo Scientific) containing 2 mL Expansion medium per well. To avoid fusion of aggregates, plates were incubated on a Celltron orbital shaker (Infors HT) at 57 rpm in an incubator at 37°C and 5% CO_2_. Plates were kept on the shaker until day 40-45.

Between days 40-45, dorsal forebrain organoids (FOs) were slicing into 300 µm air-liquid interphase ALIFO cultures using a vibratome (#VT1000S, Leica) ^3^. Then, 5-6 ALI-FOs were placed per Millicell® cell culture inserts (#PICM0RG50, Millipore) with Serum-Free Slice Culture medium beneath (SFSCM), consisting of Neurobasal medium (#21103-049, Gibco), supplemented with 1:50 (v/v) B-27 (#17504-044, Gibco), 1x (v/v) GlutaMAX (#35050-038, Thermo Scientific), 1:100 (v/v) P/S (#15140-122, Gibco) and 1:500 (v/v) Fungin (#ant-fn-2, InvivoGen). Additionally, from day 48 ALI-FOs were freshly supplemented with 0.2 µM L-ascorbic acid (AA) (#A4544, Merck), 20 ng/mL BDNF (#78133.1, SCT) and 20 ng/mL GDNF (#78139.1, SCT). After two weeks, the cell culture medium was gradually exchanged over a 3-week period from SFSCM to BrainPhys Neuronal medium (containing SM1 Supplement and N2 Supplement-A) (#05793, SCT) as well as P/S, Fungin, AA, BDNF and GDNF. The gradual shift followed this scheme: one week of 75% SFSCM and 25% BrainPhys, followed by a 50%:50% mixture in the second week, and finally a 25% SFSCM and 75% BrainPhys during the third week.

#### Sample collection

Day 40 FOs (pool of 3-5 FOs per cell line) were collected in low-binding micro tubes (#72.706.600, Sarstedt) in 500 µL 50 mM ammonium acetate (pH7.5) with PhosSTOP (#PHOSS-RO, Roche) on ice. FOs were washed once with the same solution before removing all the solution. Samples were immediately snap frozen on dry ice and stored at −70°C until further analysis. Day 80 ALI-FOs were collected from the inserts by placing a drop of PBS on top before being lifted the ALI-FO using a spatula. ALI-FOs (pool of 3-5 ALIFOs per cell line) were collected as described above.

For imaging see *Immunohistochemistry* section.

### Sample preparation for Mass Spectrometry-Based Proteomics and PTMomics

Day 40 FO and day 80 ALIFO protein and metabolite extraction, TMT labelling, PTM enrichment and high pH Reversed Phase (RP) fractionation were performed as previously described by Elmkvist et al. ^4^ with minor changes as described below. For both timepoints, samples from a total of 17 CTL and 17 SCZ-derived organoids were processed. Per timepoint, 2 sets of a TMTpro 18-plex (#A44520, Thermo Fisher Scientific) were used to label the 34 samples. In each TMT set, one channel was used for labelling a pooled sample consisting of all the included samples to enable quantitative comparison of samples between the different TMT sets.

In brief, PTMs were enriched from the same TMT labelled sample in the order described below. S-palmitoylated peptides (Depalm) were enriched from the precipitated SDC pellet after TMT labelling as described in ^5^. Phosphorylated peptides (Phospho), sialylated N-linked glycopeptides (Deglyco) and free cysteine-containing peptides (FreeCys) (labelled with CysPAT ^6^) were enriched using 5 µm TiO_2_ beads (#5020–75010, GL Science). Cysteine-containing peptides with reversible modifications (RmCys) were purified using Agarose S3 high-capacity acyl-rac capture beads (#AR-S3-2, NANOCS Inc.) from the TiO_2_ bead flow-through (FT). Lysine-acetylated peptides (LysAc) were enriched using Agarose Immunoaffinity beads carrying Anti-Lysine-acetyl antibodies (PTM-Scan Acetyl-Lysine Motif [Ac-K], #13416, Cell Signalling Technology) from the S3 beads FT. For proteomics analysis the FT from the incubation with Lys-Ac beads was used (Non-modified peptides).

### LC-MS analysis of metabolites

Metabolite samples were resuspended in 25 µL 0.1% FA of which 3 µL were transferred to a common quality control (QC) sample. A total of 4 µL of each sample was injected using a Vanquish Horizon UPLC (Thermo Fisher Scientific) equipped with a Zorbax Eclipse Plus C18 guard (2.1 × 50 mm and 1.8 µm particle size, Agilent Technologies) and an analytical column (2.1 × 150 mm and 1.8 µm particle size, Agilent Technologies) kept at 40°C. The flow rate was set to 400 µl/min using eluent A (0.1% FA) and eluent B (0.1% FA, acetonitrile): 3% B from 0 to 1.5 min, 3-40% B from 1.5 to 4.5 min, 40-95% B from 4.5 to 7.5 min, 95% B from 7.5 to 10.1 min and 95 to 3% B from 10.1 to 10.5 min before equilibration for 3.5 min with the initial conditions.

The LC flow was coupled to a TimsTOF Flex (Bruker Daltonics) for analysis in positive and negative ion modes. Metabolomics samples processed in Metaboscape (v2025, Bruker Daltonics) and annotated using the MetaboBase NIST 2020 library (Bruker Daltonics), the NIST mass spectral library (National Institute of Standards and Technology), and the Mass Bank of North America (MoNA) MS/MS library. Feature tables were blank-filtered, drift-corrected, and normalized by probabilistic quotient normalization using QC samples, then merged for positive and negative ion modes using MetaboLink^7^.

#### Metabolomics data processing

Raw data was processed with MzMine (v 3.9)^8^. In brief, the following modules were used: Mass detection, ADAP chromatogram builder, ADAP chromatogram resolver, ^13^C Isotope filter, Join aligner, and Gap filling (same RT and m/z range). Compounds were annotated at Metabolomics Standards Initiative (MSI) level 3^9^ using local MS/MS spectra databases of National Institute of Standards and Technology 17 (NIST17), MoNA and all spectral libraries available from MS-DIAL (v Aug., 2022)^10^. After compound annotation, the datasets were filtered and normalized in MetaboLink^7^. Finally, the signals were auto-scaled and log2-transformed in MetaboAnalyst^11^. To adjust for batch effects from the different organoid differentiations, we applied the ComBat algorithm as implemented in the sva package in R (version 3.60.0)^12^.

### LC-MS/MS analysis of Non-modified peptides (the proteome) and phosphopeptides/Free-Cys peptides

All peptide fractions from the High pH RP fractionation of the Non-modified peptides (20 concatenated High pH RP fractions) and phosphopeptides/FreeCys-peptides (30 concatenated High pH RP fractions) were resolubilized in 0.1% formic acid (FA) and loaded onto a Vanquish NEO system (Thermo Scientific) coupled to an Orbitrap Exploris 480 (Thermo Scientific). The peptides were loaded onto a two-column system with a 5 mm x 300 µm Acclaim™ PepMap™ 100 C18 HPLC trap column (5 µm, Thermo Scientific) using the Neo LC system. The analytical column was an in-house packed 22 cm × 100 µm ID Reprosil-Pur C18-AQ RP analytic column (1.9µm; Dr. Maisch GmbH, Germany).

The peptides were eluted from the trap column with an increasing Acetonitrile (ACN) gradient from 2% buffer B (Buffer A: 0.1% Formic Acid (FA); Buffer B: 95% ACN, 0.1% FA) to 25% B for 100 min, from 25% B to 50 % B for 20 min and finally from 50% B to 100% B for 1 min. For all peptide fractions a full MS1 was performed in the mass range of 350–1500 in the Orbitrap with a resolution of 120.000 full-width half-maximum (FWHM), a maximum ion injection time of 50 ms and a normalised automatic gain control (AGC) target of 300%. For the non-modified peptide fractions MS2 Fragmentation was performed using the Turbo-TMTpro settings at the Orbitrap Exploris 480 at high resolution (30,000 FWHM) with a normalised AGC target of 500% and a maximum injection time of 54 ms, using an isolation window of 0.7 m/z, HCD Normalized Collision Energy (NCE) of 33% and 20 sec. dynamic exclusion time. For the phosphopeptide/Free Cys-peptide fractions MS2 Fragmentation was performed using high resolution (45,000 FWHM) with a normalised AGC target of 300% and a maximum injection time of 86 ms using an isolation window of 0.7 m/z, HCD NCE of 33% and 20 sec. dynamic exclusion time.

### LC-MS/MS analysis of reversibly modified Cysteine containing peptides

Reversibly modified cysteine containing peptides (RmCys) were fractionation into 12 concatenated high pH RP fractions. All fractions were resolubilized in 0.1% FA and loaded onto a Vanquish NEO system (Thermo Scientific) coupled to an Orbitrap Exploris 480 (Thermo Scientific). The peptides were loaded onto a two-column system with a 5 mm x 300 µm Acclaim™ PepMap™ 100 C18 HPLC trap column (5 µm, Thermo Scientific) using the Neo LC system. The analytical column was an in-house packed 22 cm × 100 µm ID Reprosil-Pur C18-AQ RP analytic column (1.9µm; Dr. Maisch GmbH, Germany).

The analysis of the RmCys from day 40 was performed as described for the non-modified peptides above using TurboTMTpro settings on the Orbitrap Exploris 480, except for using a 90 min gradient (2-25%B in 70 min, 25-50%B in 20 min and 50%B-100%B in 1 min). The analysis of the RmCys fractions from day 80 was, due to more material, analyzed using a 120 min gradient similar to the phosphopeptide/FreeCys-peptide analysis described above.

### LC-MS/MS analysis of Lys-Ac peptides and N-linked deglycopeptides

The immune-purified Lys-Ac peptides and the deglycosylated formerly sialylated N-linked glycopeptides from the TiO_2_ enrichment were all resolubilized in A-buffer and loaded onto a 2 cm trap column (Reprosil 3 µm, Dr. Maisch, Ammerbuch-Entringen, Germany) using a nano-EASY LC (Thermo Fisher Scientific) coupled to an Orbitrap Exploris 480 mass spectrometer (Thermo Fisher Scientific) or an Orbitrap Astral (Thermo Fisher Scientific). The deglycopeptides were eluted using a 120 min gradient (2-25%B for 100 min, 25-50%B for 20 min and 50-100%B for 1 min) onto a 20 cm 75 μm ID analytical column containing C18 material (Reprosil 1.9 µm, Dr. Maisch, Ammerbuch-Entringen, Germany). A full MS1 was performed in the mass range of 350–1500 in the Orbitrap with a resolution of 120.000 FWHM, a maximum ion injection time of 50 ms and a normalised automatic gain control (AGC) target of 300%. For the deglycopeptides MS2 Fragmentation was performed using high resolution (50,000 FWHM) with a normalised AGC target of 300% and a maximum injection time of 100 ms using an isolation window of 0.7 m/z, HCD NCE of 33% and 20 sec. dynamic exclusion time.

The Lys-Ac peptides from day 40 was analyzed like the deglycopeptides described above. The Lys-Ac peptides from day 80 was analysed using an2 column EASY-LC coupled to an Orbitrap Astral, to increase the coverage. The Lys-Ac peptides were eluted using a 90 min gradient (2-5%B for 1 min, 5-28%B for 70 min, 28-50%B for 20 min, 50-70%B for 6 min and 70-100%B for 6 min). A full MS1 was performed in the mass range of 300–1500 in the Orbitrap with a resolution of 240.000 FWHM, a maximum ion injection time of 10 ms and a normalised automatic gain control (AGC) target of 250%. MS2 Fragmentation was performed in the Astral analyzer with a normalised AGC target of 300% and a maximum injection time of 20 ms using an isolation window of 0.7 m/z, HCD NCE of 33% and 10 sec. dynamic exclusion time.

### LC-MS/MS analysis of De-palmitoylated peptides

The de-plamitoylated peptides were fractionation into 12 concatenated high pH RP fractions. All fractions were resolubilized in 0.1% FA and loaded onto a 2 cm trap column (Reprosil 3 µm, Dr. Maisch, Ammerbuch-Entringen, Germany) using a nano-EASY LC (Thermo Fisher Scientific) coupled to an Orbitrap Exploris 480 mass spectrometer (Thermo Fisher Scientific). The de-palmitoylated peptides were sepated onto a 20 cm 75 μm ID analytical column containing C18 material (Reprosil 1.9 µm, Dr. Maisch, Ammerbuch-Entringen, Germany) using a 60 min gradient (2-28%B for 50 min, 28-50%B for 10 min and 50-100%B for 1 min) for day 40 and due to increasing amount a 120 min gradient (2-25%B for 100 min, 25-50%B for 20 min and 50-100%B) for day 80. For all samples MS1 was performed in the mass range of 350–1500 in the Orbitrap with a resolution of 120.000 FWHM, a maximum ion injection time of 50 ms and a normalised automatic gain control (AGC) target of 300%. MS2 Fragmentation was performed using high resolution (45,000 FWHM) with a normalised AGC target of 500% and a maximum injection time of 100 ms using an isolation window of 0.7 m/z, HCD NCE of 33% and 20 sec. dynamic exclusion time.

### Data search in Proteome Discoverer

All LC-MSMS data from TMT labeled peptides and modified peptides were searched in the Proteome Discoverer (PD) program (version 2.5.0.400) using the SEQUEST HT database search algorithm with the Percolator program for calculating false discovery rates. all searches were performed using the Uniprot Human annotated protein fasta database (version 26.02.2025; 20574 sequences). The search criteria were as follows; Enzyme: Trypsin (Full), Max. Missed cleavages sites 2 (for lysine-acetylated peptides 4), Precursor Mass Tolerance 10 ppm, Fragment Mass tolerance 0.05 Da, TMTpro fixed modification on N-terminals and Lysine (except for the analysis of lysine-acetylated peptides were TMTpro(K) was set as variable). For database searching of the phosphopeptides, Free Cysteine peptides (CysPAT) and deglycopeptides the variable modifications were set to: phospho (S/T/Y), SIA (C)(CysPAT modification), Carbamidomethyl (C) and deamidation (N). For lysine-acetylated peptides the variable modifications were Acetyl (K) and TMTpro (K). For the depalmitoylated peptides the variable modifications were Carbamidomethyl (C) and SIA (C)(CysPAT modification). All peptide identifications were filtered for 1% FDR using the Percolator program in PD.

#### Proteomics data analysis

Scaled non-modified protein abundances (protein level) or PTM peptide abundances (PTM level) were log_2_-transformed and only considered for subsequent analysis if quantified across both TMT 18-plexes.

Statistical analysis comparing SCZ vs CTL was performed in Proteome Discoverer (v2.5) with the in-built t-test (background based)^13^. Proteins/peptides that fulfilled the following criteria: absolute log_2_ fold change (SCZ/CTL) ≥ 0.263, adjusted p-value ≤ 0.05, and abundances (grouped) CVs ≤ 60%, were considered significantly regulated in SCZ compared to CTL.

#### SCZ-associated protein-coding genes

For each protein identified in non-modified protein and/or PTM peptide datasets, the protein was assigned as ‘SCZ-associated’ if listed with a target-disease association for ‘schizophrenia’ in the Open Targets Platform ^14^.

We selected 18 protein-coding genes associated with SCZ based on previous GWAS studies (*BCL11B, CACNA1C, CACNA1G, CUL1, FAM120A, FOXP1, GRIA3, GRIN2A, HERC1, MYT1L, RB1CC1, RERE, SETD1A, SLC39A8, SP4, STAG1, TRIO, XPO7*) ^15,16^. For each protein, the average log_2_ fold change (SCZ/CTL) of non-modified protein abundance and PTM level peptide abundances was visualized as a heatmap using the ‘ggplot2’ package in R Studio^17^. Proteins that exhibited increased/decreased levels in SCZ/CTL at the protein/PTM level were highlighted (** absolute log_2_ fold change (SCZ/CTL) ≥ 0.263, adjusted p-value ≤ 0.05, and abundances (grouped) CVs ≤ 60%, * absolute log_2_ fold change (SCZ/CTL) ≥ 0.263, adjusted p-value ≤ 0.05).

#### Pathway enrichment analysis and prediction of causal regulators

Non-modified protein abundance and PTM level data were analyzed using the QIAGEN Ingenuity Pathway Analysis (IPA) software (QIAGEN Inc.)^18^. The data was analyzed using the ‘core analysis’ functionality with the following biological filters: absolute log_2_ fold change (SCZ/CTL) ≥ 0.263, adjusted p-value ≤ 0.05. Proteins with altered abundance at the protein and/or PTM level were analyzed for pathway enrichment using a one-sided Fisher’s exact test with Benjamini-Hochberg correction for multiple testing and visualized as heatmaps using the ‘ggplot2’^17^ package in R Studio (• indicate adjusted p-value < 0.05). Potential upstream molecules/regulators of the significantly altered proteins were identified using the causal analysis function in IPA, calculating an activation Z-score and p-value of overlap (one-sided Fisher’s exact test with Benjamini-Hochberg correction) and visualized as dotplots using the ‘ggplot2’ package^17^.

#### Gene Ontology enrichment analysis

Enrichment analysis of Monarch Phenotype, KEGG pathways, and Gene Ontology (GO) terms – including Biological Process, Molecular Function, and Cellular Component – was performed on significantly regulated non-modified proteins (up or down separately) and proteins exhibiting regulation (up/down) in any post-translational modification (PTM). Analysis utilized the Cytoscape (v. 3.10.3) (Cytoscape Consortium, https://cytoscape.org/) StringApp app^19^, applying a redundancy cutoff of 0.7 to reduce term overlap. Keywords were assigned to each enriched term based on term wording, and selected summaries were generated considering keyword assignment, representation across non-modified and PTM datasets, term specificity, and average false discovery rate (FDR) for the term. Dotplots and pie charts were made with R (v4.3.0, 2023-04-21 ucrt) and RStudio (v2023.6.0.421) applying the R-packages ggplot2 (v3.5.2)^17^ and dplyr (v1.1.4)^20^.

#### Kinase enrichment analysis

The phosphopeptide data was used to predict upstream serine/threonine kinase activities employing the ‘kinase-library’ package in Python and the experimental kinase-substrate library by Johnson et al.^21^. Kinase-substrate relationships were predicted for serine/threonine kinases (percentile-rank, threshold: 15) and kinase enrichment was performed for phosphorylation sites with significant up- (log_2_ fold change (SCZ/CTL) ≥ 0.263, adjusted p-value ≤ 0.05, and abundances (grouped) CVs ≤ 60%) or downregulation (log_2_ fold change (SCZ/CTL) ≤ −0.263, adjusted p-value ≤ 0.05, and abundances (grouped) CVs ≤ 60%) when comparing SCZ and CTL. Non-regulated phosphorylation sites were used as background. Kinase enrichment was calculated using a one-sided Fisher’s exact test with Benjamini-Hochberg correction for multiple testing. Kinases with an absolute log_2_(Frequency factor) score ≥ 0.5 and adjusted p-value ≤ 0.05 were considered as having increased/decreased activity in SCZ compared to CTL. Kinase enrichment results were plotted as a Volcano plot using the ‘ggplot2’ package in R Studio^17^.

#### Data visualization

Unless otherwise specified above, bar plots, circle diagrams and boxplots were prepared using GraphPad Prism (v10.6.1) or R. Overlap between the number of proteins with regulations at the protein and PTM level was visualized as euler plots using the ‘eulerr’ and ‘ggplot2’ packages in R Studio^17^. Differentially expressed proteins were visualized using Volcano plots (using the ‘ggplot2’ package in R Studio) and enrichment analyses were performed using clusterProfiler and enrichplot with all identified proteins as background^22,23^, and results were corrected for multiple comparisons using the Benjamini–Hochberg method.

### Data Independent Acquisition (DIA) LC–MS/MS for Label-Free Protein Quantification

All peptide samples from day 40 and day 80 was subjected to Data Independent Acquisition using a TIMS-TOF HT instrument (Bruker Daltonics). A Vanquish Neo LC pumps (Thermo scientific) were used to deliver peptide samples for analysis to Bruker timsTOF HT instrument. The LC setup: precolumn – PEPMAP NEO C18 5µm 300µm x 5mm trap cartridge. Trapped peptides (200ng loaded) were eluted and separated on an analytical column (PepSep Fifteen series (C18, 15cm × 100µm, 3µm resin size, Bruker)) in a 42 min gradient of increasing concentration of solvent B (95% ACN, 0.1% FA (solvent A – 0.1% FA). Peptides were analyzed in dia-PASEF mode with the following settings: MS scan 100-1700 m/z, positive polarity; TIMS: 1/K0 Start – End 0.6 – 1.6 V×s/cm^2^, ramp time 100ms. dia-PASEF: cycle time 1.59 sec, 14 MS/MS ramps, 29 MS/MS windows, mass range: 340.3 – 1210.3 m/z, mobility range 0.66 – 1.44 1/K0.

#### DIA raw data search for label-free quantification

The raw data were searched by Spectronaut software (v20) against human protein database (fasta file, v2025) utilizing chiefly its standard settings with modifications as follows. Cleavage: trypsin/P, specific, peptide length 7-52 AA, max 2 missed cleavages; modifications: fixed carbamidomethylation of Cys, variable oxidation of Met and acetylation of protein N-term. Identification: directDIA+ (Deep), Protein Qvalue Cutoff (Experiment): 0.01. Quantification: MS Level: MS2, type “Area”, Cross-Run Normalization applied. List of identified and quantified proteins was then exported.

### Single nuclei RNA (snRNA) sample preparation and sequencing

#### Nuclei isolation and fixation

Nuclei isolation and fixation was performed on a total of 3-5 FOs (day 40) or 4-7 ALI-FOs (day 80) from 12 CTL cell lines and 12 SCZ cell lines (in total 48 samples) spanning 4 batches with equal numbers of CTL and SCZ from each batch. FOs and ALI-FOs were collected in D-PBS on ice followed by careful removal of D-PBS and snap-frozen on dry ice. Samples were stored at −70°C until further processing.

Samples were processed using the gentleMACS Dissociator (Miltenyi Biotec) in combination with C-tubes (#130-096-334, Miltenyi Biotec) to ensure efficient mechanical dissociation while preserving nuclear integrity. All steps were performed cold using pre-chilled tubes. Four samples were processed at the same time. Briefly, samples were transferred to C-tubes using 1 mL lysis buffer (Nuclei extraction buffer (#130-128-024, Miltenyi Biotec) containing 9 U/mL DNase I (M0303S, New England Biolabs) and 0.2 U/µL RNase inhibitor (#M0307L, New England Biolabs)). The tubes were mounted on the gentleMACS Dissociator placed at 4°C and subjected to a pre-programmed dissociation protocol (4C-nuclei_1 program (5 min)). Following dissociation, the homogenate was placed on ice for 5 min and thereafter passed through a MACS SmartStrainer (70 µm, #130-098-462, Miltenyi Biotech) into a 5 mL LoBind® tube (#0030122356, Eppendorf). Rinse the C-tube with another 1 mL Lysis buffer and pass over the same strainer. Apply the nuclei suspension to a MACS SmartStrainer (30 µm, #130-098-458, Miltenyi Biotech) placed on another 5 mL LoBind® tube. Depending on the amount of starting material 0.5-1 mL sample was transferred to a new 5 mL tube and centrifuged at 300×g for 10 minutes at 4°C (using swinging bucket rotor). The resulting pellet was resuspended in 450 µL Nuclei separation buffer (NSB) (Nuclei extraction buffer (Miltenyi Biotec) diluted 1:7 with PBS pH 7.2 (#20012019, Gibco) and containing 0.04% BSA (A9418, Sigma) and 2 U/µL RNase inhibitor. To purify the nuclei, 50 µL/sample anti-nucleus microbeads (#130-132-997, Miltenyi Biotech) were added, mixed by pipetting and incubated for 15 min at 4°C. During incubation, 4 LS columns (#130-042-401, Miltenyi Biotech) were prepared on a QuadroMACS separator (Miltenyi Biotech, 130-090-976) by rinsing with 3 ml NSB (wo RNase inhibitor). Samples were diluted with 2 ml NSB (wo RNase inhibitor) and applied to the LS columns. Flow-through was collected in Falcon tubes. The LS columns were washed 2x with NSB (wo RNase inhibitor) before the columns were removed from the separator and placed on a 5 mL LoBind® tubes. Nuclei were eluted by adding 1 mL NSB with RNase inhibitor to the column and immediately flush by pushing the plunger into the column. To count the nuclei, 10 µL/sample were collected and counted using a Countess II Automated cell counter (Thermo Fisher) using Trypan Blue Stain (#T10282, Thermo). Volumes were adjusted to have <1,000,000 nuclei/sample for the fixation steps. Samples were centrifuged at 400×g for 10 minutes at 4°C (swinging bucket) and the supernatant carefully aspirated. Nuclei fixation was performed using the Evercode™ Nuclei Fixation v3 kit (#ECFN3300, Parse Bioscience) according to the manufacturer’s protocol. In brief, nuclei pellets were resuspended in 187.5 µL Cell Prefix Master Mix followed by adding 62.5 µL Fixative solution and immediately mixed by pipetting exactly 3 times. Samples were centrifuged at 400×g for 10 minutes at 4°C (swinging bucket) and the supernatant removed subsequently. The nuclei pellets were resuspended in 100 µL Cell Storage Master Mix and passed over a MACS SmartStrainer (30 µm) into a 1.5 mL Low-binding tube (72.706.600, Sarstedt). For post-thaw nuclei count, 10 µL/sample were collected in 0.5 mL Eppendorf tubes. All samples were frozen in Mr. Frosty freezing containers (5100-0001, Nalgene) at −70°C.

#### snRNA library preparation and sequencing

The 48 nuclei samples were stored at −80 °C, counted manually using a Neubauer counting chamber, and their quality was assessed. Barcoding and library preparation were performed using the PARSE Biosciences Whole Transcriptome Kit (v3, #UMWT3300) according to the manufacturer’s recommendations. A total of 8 libraries were generated, each consisting of 12000 nuclei. cDNA concentrations were quantified using the Qubit HS dsDNA Assay Kit (#Q32854, Thermo Fisher Scientific) and Qubit Fluorometer and quality was assessed using the High Sensitivity D5000 Reagents (#5067-5593, Agilent) and Agilent 4150 TapeStation. Libraries were pooled according to the expected number of nuclei per sample. The pooled libraries were quantified and sequenced using Illumina S2 Reagent Kit v1.5 (100 cycles) runs on an Illumina NovaSeq 6000 system (#20012850, Illumina), controlled by NovaSeq Control Software.

#### Data preprocessing and cell filtering

Primary data processing was performed using bcl2fastq (v2.20.0, Illumina) to generate FASTQ files, followed by split-pipe (v1.6.1, Parse Biosciences) for read alignment and count matrix generation. Reads were aligned to the Ensembl Homo sapiens GRCh38 reference genome (release 109). Each of the eight sublibraries was processed individually and subsequently merged to generate a unified count matrix per sample.

Cell filtering was carried out using the development version of CRMetrics (https://github.com/khodosevichlab/CRMetrics), which bundles standard preprocessing steps and provides visual assessment of filtered cells^24^. Scrublet was used to calculate doublet scores, and only cells below the sample-specific threshold were retained^25^. In addition, cells with a mitochondrial gene fraction > 5% or fewer than 500 or 1000 unique molecular identifiers (UMIs), depending on the sample, were excluded.

#### Sample integration and clustering

For each dataset (D40 and D80), filtered count matrices were integrated and embedded using a custom helper function wrapping Pagoda2 and Conos^26,27^. Each sample’s count matrix was independently preprocessed using Pagoda2 (with min.cells.per.gene = 3, min.transcripts.per.cell = 200), after which all samples were jointly embedded into a UMAP graph using Conos with an alignment strength of 0.2. One sample (SCZ032_D80) was excluded from downstream analysis due to low amount of detected cells, leaving 23 samples in the D80 dataset. Filtered count matrices from D40 (24 samples) and D80 (23 samples, after exclusion of SCZ032_D80) were merged into a single dataset of 47 samples and jointly embedded using the same Conos/Pagoda2 pipeline described above. An alignment strength of 0.2 was applied, consistent with the individual timepoint analyses. Leiden community detection was run at a resolution of 2.5, yielding 16 clusters. Cell type annotation was performed based known marker genes from the literature. Based on marker gene expression overlap, four clusters were combined into one cortical neuron (CorNeuron) cluster and two into one outer radial glia (oRG) cluster, resulting in 12 final clusters. Further analyses were performed using Seurat in R and Trailmaker^TM^ (Parse Biosciences)^28,29^. Cluster-specific marker genes were identified by comparing cells of each cluster to all other cells using the presto package implementation of the Wilcoxon rank-sum test.

#### Differential gene expression, enrichment analysis and pseudotime analysis

Differentially expressed genes (DEGs) between SCZ and CTL at day 40 and day 80 were identified using a pseudobulk limma-voom workflow in Trailmaker. This was performed both at the group level and cell cluster level. Significance was determined at an adjusted p-value threshold of 0.05 and a log2 fold change cutoff of 0.5. Enrichment analyses were subsequently performed using clusterProfiler and enrichplot^22,23^, and results were corrected for multiple comparisons using the Benjamini–Hochberg method. The top 20 regulated biological process (BP) gene ontology (GO) terms were visualized as clustered heatmaps to reduce redundancy among similar terms using gplots^30^. The top four up- and downregulated BP GO terms and their connected DEGs were visualized in cnetplots.

Pseudotime analysis was performed using Monocle3 by subsetting the SCZ and CTL data and converting their integrated Seurat object^31^. UMAP coordinates and clusters were assigned from Seurat and trajectory analysis performed not considering cluster partitions. Cells were ordered by denoting the cluster of ventricular radial glia (vRG) as starting points.

#### Sex-stratified differential expression analysis

To investigate sex-specific transcriptional differences in SCZ, sex-stratified pseudobulk DEG analyses were performed for both timepoints (D40 and D80). Pseudobulk limma-voom tests were applied as described above, comparing SCZ to CTL within female and male donors separately.

### Immunocytochemistry - iPSCs

Coverslips were fixated in 4% paraformaldehyde (PFA, #28908, Thermo Fisher Scientific) in PBS (#14190-144, Thermo Fisher Scientific) for 20 min at RT and washed twice in PBS for 15 min before permeabilization and blocking of unspecific antibody binding in blocking buffer (PBS/0.1% Triton X-100 (#T-9284, Sigma)/5% goat serum (#16210-064, Thermo Fisher Scientific)) for 1 hour at RT. Primary antibodies were diluted in blocking buffer and incubated ON at 4°C. Primary antibodies and dilutions are listed in Table S3. After primary antibody incubation, cultures were washed three times in wash buffer (PBS/0.1% Triton X-100) for 15 min at RT and incubated with appropriate fluorophore-conjugated secondary antibodies and nuclei counterstained with DAPI (NucBlue, #R37606, Thermo Fischer Scientific, 1:1000) diluted in blocking buffer, ON at 4°C. Secondary antibodies and dilutions are listed in Table S3. After secondary antibody incubation, cultures were washed three times in PBS for 10 min and coverslips mounted on glass slides with ProLong Diamond antifade mounting medium (#P36961, Thermo Fischer Scientific).

### Immunohistochemistry - FOs and ALIFOs

Whole organoids (day 20 samples) were fixated in 4% PFA in PBS for 20 min at RT, while ALIFOs (day 40 and day 80 samples) were fixated in 4% PFA in PBS for 1 hour at RT, and subsequently soaked in 30% sucrose in PBS at 4°C for at least 1 day. Three organoids or ALIFOs per cell line were washed twice in washing buffer (PBS/0.3% saponin (#84510, Sigma) or PBS/0.25% Triton X-100) for 15 min before permeabilization and blocking of unspecific antibody binding in either PBS/0.3% saponin/5% goat serum or PBS/0.25% Triton X-100/5% goat serum for 1 hour at RT. Samples were permeabilized with saponin when detecting membrane proteins and neuronal morphology, and with Triton X-100 when detecting nuclear proteins. Primary antibodies were diluted in blocking buffer and incubated for 2 days at 4°C with gentle shaking. Primary antibodies and dilutions are listed in Table S3. After primary antibody incubation, samples were washed three times in washing buffer for 1 hour at RT with gentle shaking, before incubation with appropriate fluorophore-conjugated secondary antibodies and nuclei counterstained with DAPI diluted in blocking buffer ON at 4°C with gentle shaking. Secondary antibodies and dilutions are listed in Table S3. After secondary antibody incubation, samples were washed three times in PBS for 30 min at RT and cleared using RapiClear 1.47 solution (#RC147002, SunJin Lab) for at least 1 day at RT with gentle shaking. When sufficiently cleared, samples were mounted in RapiClear solution on glass slides within an 0.3 mm deep iSpacer double layer ring (#IS317, SunJin Lab).

### Image Acquisition

Image acquisition was performed at the Danish Molecular Biomedical Imaging Center (DaMBIC, University of Southern Denmark). Confocal images were acquired on the AX Confocal/Multiphoton microscope (Nikon) at X10, X20 or X60 magnification depending on the staining. 4 randomly chosen fields/sample were imaged with identical settings for all samples. Detailed imaging information for each staining are listed in Table S4.

Bright field images of live organoids during differentiation were acquired with an EVOS brightfield microscope (Invitrogen) at X4 (early timepoints) or X2 (later timepoints) magnification.

### Image analysis

Image analyses were performed in ImageJ/Fiji (NIH) using custom-written macros, depending on the type of image analysis, to ensure standardized image processing. For stainings acquired as z-stacks, maximum intensity projections were generated prior to analysis to ensure that marker-positive structures throughout the imaged volume were represented in a single image plane.

For cell type quantification of nuclear staining total cell counts were quantified from the DAPI staining. DAPI images were background-subtracted using the rolling ball algorithm, contrast was enhanced, followed by Gaussian blurring to reduce high-frequency noise. Nuclei were segmented using the “Find Maxima” function (output: segmented particles) and thresholding performed to create a binary mask. The specific cell type staining was also background-subtracted, followed by segmentation using the “Find Maxima” function and thresholding performed to create a binary mask. Quantification was performed on the binary mask using the “Analyze Particles” function and cell type counts normalized to total cell count from the DAPI quantification.

Cell type quantification of whole cell staining was estimated from total staining intensity by measuring mean grey values using the “Measure” function and normalizing to mean grey values of the DAPI staining.

For synapse quantifications, BTUB^+^-regions were segmented to define neuronal areas. Synapse staining was background-subtracted, contrast-enhanced, and thresholding performed to create binary masks. Synaptic puncta overlapping with BTUB^+^-regions were selected, and the “Analyze Particles” function was applied to quantify the number of presynaptic, postsynaptic, and colocalized puncta. Synapse density was normalized to the BTUB^+^-area.

Organoid growth curves were analysed from bright field images of live organoids. Images were converted to 8-bit grayscale and subjected to Gaussian blur to reduce noise. Thresholding was performed to create a binary mask, and watershed segmentation was applied to separate touching organoids. The area of the organoids was measured on the binary mask using the “Analyze Particles” function. At least 10 organoids were analysed per cell line for each timepoint.

Neuronal migration was analysed on doublecortin (DCX) stainings. Identical brightness and contrast settings were applied to all images and kept constant throughout the analysis. Both staining localization and radial migration was assessed. As the analysis was performed manually, images were blinded prior to quantification. For each rosette, three regions of interest (ROIs) were manually defined using the “Polygon Selection tool”: (1) the rosette lumen, (2) the entire rosette, and (3) the peripheral area surrounding the rosette up to the border of neighbouring rosettes. Integrated density was measured for each ROI. Peripheral DCX intensity was calculated by subtracting the integrated density of the full rosette from that of the surrounding area, representing migrated neurons. DCX intensity within the lumen was used as a measure of non-migrated neurons. Only rosettes with a clearly visible lumen, a fully visible surrounding area, and no contact with the organoid edge were included.

### Western blotting

Protein samples were as described for DDA proteomics above. For each sample, 10 µg protein was separated using Bolt 4-12% Bis-Tris Plus Gels (#NW04125, Thermo Fisher Scientific) with samples loaded in NuPage LDS Sample buffer (#NP0007, Thermo Fisher Scientific) with NuPage Sample Reducing agent (#NP0004, Thermo Fisher Scientific) and PageRuler Plus pre-stained protein standard (#26620, Thermo Fisher Scientific) for size reference. Proteins were transferred to activated PVDF membranes (#IPVH00010, Millipore) using the Trans-Blot® Turbo™ Transfer System (BioRad) and membranes were blocked with 3% Bovine serum albumin (BSA, #A3059, Sigma-Aldrich) in TBS-Tween20 (TBST) buffer for 1 h and incubated overnight at 4°C in 3% BSA/TBST with anti-rabbit phospho-YAP (Ser127) (#PA517481, Invitrogen) 1:1000. Anti-mouse β-actin HRP-linked (#ab49900, Abcam) 1:50,000 was used as loading control. Following 3 x wash in TBST, the membranes were incubated for 1 h at RT in TBST with the secondary anti-rabbit IgG, HRP-linked (Cell Signalling, #7074) 1:10,000. Following 3 x wash in TBST, the membranes were visualized with Immobilon Forte Western HRP Substrate (# WBLUF0100, Millipore) using an Amersham 680 Imager (GE Healthcare).

For quantification, the optical density of each band was quantified using Image Lab software (Bio-Rad).

## Acknowledgements

The project was funded by the Lundbeck Foundation under the DEVELOPNOID project (grant no. R336+2020-1113).

We acknowledge Laura Wolbeck, Irina Korshunova and Eman Ahmad Mouhammad and the Neuro Single Cell Platform - NeuSiC at BRIC, the University of Copenhagen, funded by the Lundbeck Foundation (grant no. R433-2023-1585), for technical and computational expertise and support.

Image acquisition was performed at the Danish Molecular Biomedical Imaging Center (DaMBIC, University of Southern Denmark), supported by the Novo Nordisk Foundation (NNF) (grant agreement no. NNF18SA0032928).

This project was supported by a generous grant from the Danish Agency of Higher Education and Science to establish the PLATO research infrastructure: Danish National Mass Spectrometry Platform for Proteomics and Biomolecular Imaging (grant no. 5229-00012B, www.sdu.dk/PLATO).

The Villum Center for Bioanalytical Sciences at the University of Southern Denmark is acknowledged for access to state-of-the-art mass spectrometry instrumentation.

## Author contributions

H.B., S.I.S, P.J. and M.R.L. conceived and designed the study. H.B., S.I.S., M.S.Ø., S.B.E., P.J., F.A.M., A.W.M., J.F.H., M.R., L.C., L.A.J. and A.N. performed experiments and acquired data. H.B., S.I.S., M.S.Ø., S.B.E., P.J, J.F.H. and M.R.L. analysed and interpreted the data. E.B. and M.E.B. contributed with clinical expertise, patient recruitment, and clinical characterization. J.B. provided imaging support. N.J.F. contributed expertise in metabolomics and data interpretation. M.L. and P.R. assisted with methodology and supervision. H.B. and S.I.S drafted the manuscript. M.R.L. supervised the study and acquired funding. All authors contributed to data interpretation and provided critical feedback in writing the paper.

## Main references

1 Hilker, R. et al. Heritability of Schizophrenia and Schizophrenia Spectrum Based on the Nationwide Danish Twin Register. Biol Psychiatry 83, 492–498 (2018). 10.1016/j.biopsych.2017.08.017

2 Trubetskoy, V. et al. Mapping genomic loci implicates genes and synaptic biology in schizophrenia. Nature 604, 502–508 (2022). 10.1038/s41586-022-04434-5

3 Sullivan, P. F., Yao, S. & Hjerling-Leffler, J. Schizophrenia genomics: genetic complexity and functional insights. Nat Rev Neurosci 25, 611–624 (2024). 10.1038/s41583-024-00837-7

4 Sullivan, P. F. & Geschwind, D. H. Defining the Genetic, Genomic, Cellular, and Diagnostic Architectures of Psychiatric Disorders. Cell 177, 162–183 (2019). 10.1016/j.cell.2019.01.015

5 Birnbaum, R. & Weinberger, D. R. Genetic insights into the neurodevelopmental origins of schizophrenia. Nat Rev Neurosci 18, 727–740 (2017). 10.1038/nrn.2017.125

6 Pasca, S. P. The rise of three-dimensional human brain cultures. Nature 553, 437–445 (2018). 10.1038/nature25032

7 Pasca, A. M. et al. Functional cortical neurons and astrocytes from human pluripotent stem cells in 3D culture. Nat Methods 12, 671–678 (2015). 10.1038/nmeth.3415

8 Gordon, A. et al. Long-term maturation of human cortical organoids matches key early postnatal transitions. Nat Neurosci 24, 331–342 (2021). 10.1038/s41593-021-00802-y

9 Brennand, K. J. et al. Modelling schizophrenia using human induced pluripotent stem cells. Nature 473, 221–225 (2011). 10.1038/nature09915

10 Kathuria, A. et al. Transcriptomic Landscape and Functional Characterization of Induced Pluripotent Stem Cell-Derived Cerebral Organoids in Schizophrenia. JAMA Psychiatry 77, 745–754 (2020). 10.1001/jamapsychiatry.2020.0196

11 Sawada, T. et al. Developmental excitation-inhibition imbalance underlying psychoses revealed by single-cell analyses of discordant twins-derived cerebral organoids. Mol Psychiatry 25, 2695–2711 (2020). 10.1038/s41380-020-0844-z

12 Rasanen, N. et al. miRNA profiling of hiPSC-derived neurons from monozygotic twins discordant for schizophrenia. Schizophrenia (Heidelb) 11, 21 (2025). 10.1038/s41537-025-00573-6

13 Liu, Y., Beyer, A. & Aebersold, R. On the Dependency of Cellular Protein Levels on mRNA Abundance. Cell 165, 535–550 (2016). 10.1016/j.cell.2016.03.014

14 Chandrasekaran, A. et al. A protein-centric view of in vitro biological model systems for schizophrenia. Stem Cells 39, 1569–1578 (2021). 10.1002/stem.3447

15 Bludau, I. & Aebersold, R. Proteomic and interactomic insights into the molecular basis of cell functional diversity. Nat Rev Mol Cell Biol 21, 327–340 (2020). 10.1038/s41580-020-0231-2

16 Giandomenico, S. L. et al. Cerebral organoids at the air-liquid interface generate diverse nerve tracts with functional output. Nat Neurosci 22, 669–679 (2019). 10.1038/s41593-019-0350-2

17 Bestman, J. E., Huang, L. C., Lee-Osbourne, J., Cheung, P. & Cline, H. T. An in vivo screen to identify candidate neurogenic genes in the developing Xenopus visual system. Dev Biol 408, 269–291 (2015). 10.1016/j.ydbio.2015.03.010

18 Jeon, C. Y. et al. p190RhoGAP and Rap-dependent RhoGAP (ARAP3) inactivate RhoA in response to nerve growth factor leading to neurite outgrowth from PC12 cells. Exp Mol Med 42, 335–344 (2010). 10.3858/emm.2010.42.5.035

19 Gordon, A. et al. Developmental convergence and divergence in human stem cell models of autism. Nature 651, 707–719 (2026). 10.1038/s41586-025-10047-5

20 Rao, S. B. et al. Aberrant pace of cortical neuron development in brain organoids from patients with 22q11.2 deletion syndrome-associated schizophrenia. Nat Commun 16, 6986 (2025). 10.1038/s41467-025-62187-x

21 Impagnatiello, F. et al. A decrease of reelin expression as a putative vulnerability factor in schizophrenia. Proc Natl Acad Sci U S A 95, 15718–15723 (1998). 10.1073/pnas.95.26.15718

22 Ovadia, G. & Shifman, S. The genetic variation of RELN expression in schizophrenia and bipolar disorder. PLoS One 6, e19955 (2011). 10.1371/journal.pone.0019955

23 Vilchez-Acosta, A. et al. Specific contribution of Reelin expressed by Cajal-Retzius cells or GABAergic interneurons to cortical lamination. Proc Natl Acad Sci U S A 119, e2120079119 (2022). 10.1073/pnas.2120079119

24 Singh, T. et al. Rare coding variants in ten genes confer substantial risk for schizophrenia. Nature 604, 509–516 (2022). 10.1038/s41586-022-04556-w

25 Vogel, W. K., Gafken, P. R., Leid, M. & Filtz, T. M. Kinetic analysis of BCL11B multisite phosphorylation-dephosphorylation and coupled sumoylation in primary thymocytes by multiple reaction monitoring mass spectroscopy. J Proteome Res 13, 5860–5868 (2014). 10.1021/pr5007697

26 Sarikas, A., Hartmann, T. & Pan, Z. Q. The cullin protein family. Genome Biol 12, 220 (2011). 10.1186/gb-2011-12-4-220

27 Besold, A. N. & Michel, S. L. Neural Zinc Finger Factor/Myelin Transcription Factor Proteins: Metal Binding, Fold, and Function. Biochemistry 54, 4443–4452 (2015). 10.1021/bi501371a

28 Notaras, M., Lodhi, A., Fang, H., Greening, D. & Colak, D. The proteomic architecture of schizophrenia iPSC-derived cerebral organoids reveals alterations in GWAS and neuronal development factors. Transl Psychiatry 11, 541 (2021). 10.1038/s41398-021-01664-5

29 Bement, W. M., Goryachev, A. B., Miller, A. L. & von Dassow, G. Patterning of the cell cortex by Rho GTPases. Nat Rev Mol Cell Biol 25, 290–308 (2024). 10.1038/s41580-023-00682-z

30 Unzueta-Larrinaga, P. et al. Extracellular matrix dysfunction and synaptic alterations in schizophrenia. Mol Psychiatry 31, 396–406 (2026). 10.1038/s41380-025-03154-2

31 Gomez, A. M., Traunmuller, L. & Scheiffele, P. Neurexins: molecular codes for shaping neuronal synapses. Nat Rev Neurosci 22, 137–151 (2021). 10.1038/s41583-020-00415-7

32 Li, Y. et al. Protein Kinase C Family: Structures, Biological Functions, Diseases, and Pharmaceutical Interventions. MedComm (2020) 6, e70474 (2025). 10.1002/mco2.70474

33 Wayman, G. A., Lee, Y. S., Tokumitsu, H., Silva, A. J. & Soderling, T. R. Calmodulin-kinases: modulators of neuronal development and plasticity. Neuron 59, 914–931 (2008). 10.1016/j.neuron.2008.08.021

34 Hu, C., Feng, P., Yang, Q. & Xiao, L. Clinical and Neurobiological Aspects of TAO Kinase Family in Neurodevelopmental Disorders. Front Mol Neurosci 14, 655037 (2021). 10.3389/fnmol.2021.655037

35 Frank, C. L. & Tsai, L. H. Alternative functions of core cell cycle regulators in neuronal migration, neuronal maturation, and synaptic plasticity. Neuron 62, 312–326 (2009). 10.1016/j.neuron.2009.03.029

36 Onishi, K. et al. LRRK2 mediates axon development by regulating Frizzled3 phosphorylation and growth cone-growth cone communication. Proc Natl Acad Sci U S A 117, 18037–18048 (2020). 10.1073/pnas.1921878117

37 Byeon, S. & Yadav, S. Pleiotropic functions of TAO kinases and their dysregulation in neurological disorders. Sci Signal 17, eadg0876 (2024). 10.1126/scisignal.adg0876

38 Takemoto-Kimura, S. et al. Calmodulin kinases: essential regulators in health and disease. J Neurochem 141, 808–818 (2017). 10.1111/jnc.14020

39 Malhotra, D. et al. High frequencies of de novo CNVs in bipolar disorder and schizophrenia. Neuron 72, 951–963 (2011). 10.1016/j.neuron.2011.11.007

40 van Woerden, G. M. et al. TAOK1 is associated with neurodevelopmental disorder and essential for neuronal maturation and cortical development. Hum Mutat 42, 445–459 (2021). 10.1002/humu.24176

41 Kaiser, J. et al. Convergence on CaMK4: A Key Modulator of Autism-Associated Signaling Pathways in Neurons. Biol Psychiatry 97, 439–449 (2025). 10.1016/j.biopsych.2024.10.012

42 Bentea, E. et al. Kinase network dysregulation in a human induced pluripotent stem cell model of DISC1 schizophrenia. Mol Omics 15, 173–188 (2019). 10.1039/c8mo00173a

43 Sebastien, M., Paquette, A. L., Prowse, E. N. P., Hendricks, A. G. & Brouhard, G. J. Doublecortin restricts neuronal branching by regulating tubulin polyglutamylation. Nat Commun 16, 1749 (2025). 10.1038/s41467-025-56951-2

44 Zhuo, C., Hou, W., Tian, H., Wang, L. & Li, R. Lipidomics of the brain, retina, and biofluids: from the biological landscape to potential clinical application in schizophrenia. Transl Psychiatry 10, 391 (2020). 10.1038/s41398-020-01080-1

45 Reay, W. R. & Cairns, M. J. The role of the retinoids in schizophrenia: genomic and clinical perspectives. Mol Psychiatry 25, 706–718 (2020). 10.1038/s41380-019-0566-2

46 Lavado, A. et al. YAP/TAZ maintain the proliferative capacity and structural organization of radial glial cells during brain development. Dev Biol 480, 39–49 (2021). 10.1016/j.ydbio.2021.08.010

47 Lian, K. et al. Hub genes, a diagnostic model, and immune infiltration based on ferroptosis-linked genes in schizophrenia. IBRO Neurosci Rep 16, 317–328 (2024). 10.1016/j.ibneur.2024.01.007

48 Steele-Perkins, G. et al. The transcription factor gene Nfib is essential for both lung maturation and brain development. Mol Cell Biol 25, 685–698 (2005). 10.1128/MCB.25.2.685-698.2005

49 Brenes, A. J. et al. Erosion of human X chromosome inactivation causes major remodeling of the iPSC proteome. Cell Rep 35, 109032 (2021). 10.1016/j.celrep.2021.109032

50 Shelly, M. et al. Semaphorin3A regulates neuronal polarization by suppressing axon formation and promoting dendrite growth. Neuron 71, 433–446 (2011). 10.1016/j.neuron.2011.06.041

51 (!!! INVALID CITATION !!! 51).

52 Tiihonen, J. et al. Sex-specific transcriptional and proteomic signatures in schizophrenia. Nat Commun 10, 3933 (2019). 10.1038/s41467-019-11797-3

53 Koopmans, F. et al. Human brain prefrontal cortex proteomics identifies compromised energy metabolism and neuronal function in Schizophrenia. Nat Commun 17 (2026). 10.1038/s41467-026-68950-y

54 Choi, J. H. et al. Phospholipase C-gamma1 is a guanine nucleotide exchange factor for dynamin-1 and enhances dynamin-1-dependent epidermal growth factor receptor endocytosis. J Cell Sci 117, 3785–3795 (2004). 10.1242/jcs.01220

55 Liu, Y., Tao, Y. M., Woo, R. S., Xiong, W. C. & Mei, L. Stimulated ErbB4 internalization is necessary for neuregulin signaling in neurons. Biochem Biophys Res Commun 354, 505–510 (2007). 10.1016/j.bbrc.2007.01.009

56 Liu, Z., Velpula, K. K. & Devireddy, L. 3-Hydroxybutyrate dehydrogenase-2 and ferritin-H synergistically regulate intracellular iron. FEBS J 281, 2410–2421 (2014). 10.1111/febs.12794

57 Godinez, A. et al. Neuroserpin, a crucial regulator for axogenesis, synaptic modelling and cell-cell interactions in the pathophysiology of neurological disease. Cell Mol Life Sci 79, 172 (2022). 10.1007/s00018-022-04185-6

58 Steffek, A. E., McCullumsmith, R. E., Haroutunian, V. & Meador-Woodruff, J. H. Cortical expression of glial fibrillary acidic protein and glutamine synthetase is decreased in schizophrenia. Schizophr Res 103, 71–82 (2008). 10.1016/j.schres.2008.04.032

59 Yang, Y. et al. Endophilin A1 drives acute structural plasticity of dendritic spines in response to Ca2+/calmodulin. J Cell Biol 220 (2021). 10.1083/jcb.202007172

60 Xu, S., Rigaux, E., Hene, D., Renard, H. F. & Thines, L. Bending the boundaries: the many facets of endophilin-As from membrane dynamics to disease. Cell Mol Life Sci 82, 339 (2025). 10.1007/s00018-025-05856-w

## Method References

1 Mohamed, F. A. et al. Dysregulation of miRNAs Drives Premature GABAergic Maturation and Early Neurodevelopmental Defects in Schizophrenia. bioRxiv, 2026.2005.2025.727574 (2026). 10.64898/2026.05.25.727574

2 Ohlenschlaeger, M. S. et al. Multi-omic analysis of guided and unguided forebrain organoids reveals differences in cellular composition and metabolic profiles. Cell Rep Methods 6, 101295 (2026). 10.1016/j.crmeth.2025.101295

3 Giandomenico, S. L. et al. Cerebral organoids at the air-liquid interface generate diverse nerve tracts with functional output. Nat Neurosci 22, 669–679 (2019). 10.1038/s41593-019-0350-2

4 Elmkvist, S. B. et al. Temporal proteomic and PTMomic atlas of cerebral organoid development. bioRxiv, 2024.2009.2003.610941 (2024). 10.1101/2024.09.03.610941

5 Jensen, P. T., Palmisano, G., Rhodes, C. J. & Larsen, M. R. Enrichment of Cysteine S-palmitoylated Peptides Using Sodium Deoxycholate Acid Precipitation. Mol Cell Proteomics 25, 101218 (2026). 10.1016/j.mcpro.2025.101218

6 Huang, H. et al. Simultaneous Enrichment of Cysteine-containing Peptides and Phosphopeptides Using a Cysteine-specific Phosphonate Adaptable Tag (CysPAT) in Combination with titanium dioxide (TiO2) Chromatography. Mol Cell Proteomics 15, 3282–3296 (2016). 10.1074/mcp.M115.054551

7 Mendes, A., Havelund, J. F., Lemvig, J., Schwammle, V. & Faergeman, N. J. MetaboLink: A web application for Streamlined Processing and Analysis of Large-Scale Untargeted Metabolomics Data. Bioinformatics 40 (2024). 10.1093/bioinformatics/btae459

8 Schmid, R. et al. Integrative analysis of multimodal mass spectrometry data in MZmine 3. Nat Biotechnol 41, 447–449 (2023). 10.1038/s41587-023-01690-2

9 Sumner, L. W. et al. Proposed minimum reporting standards for chemical analysis Chemical Analysis Working Group (CAWG) Metabolomics Standards Initiative (MSI). Metabolomics 3, 211–221 (2007). 10.1007/s11306-007-0082-2

10 Tsugawa, H. et al. MS-DIAL: data-independent MS/MS deconvolution for comprehensive metabolome analysis. Nat Methods 12, 523–526 (2015). 10.1038/nmeth.3393

11 Pang, Z. et al. MetaboAnalyst 5.0: narrowing the gap between raw spectra and functional insights. Nucleic Acids Res 49, W388–W396 (2021). 10.1093/nar/gkab382

12 Johnson, W. E., Li, C. & Rabinovic, A. Adjusting batch effects in microarray expression data using empirical Bayes methods. Biostatistics 8, 118–127 (2007). 10.1093/biostatistics/kxj037

13 Navarro, P. et al. General statistical framework for quantitative proteomics by stable isotope labeling. J Proteome Res 13, 1234–1247 (2014). 10.1021/pr4006958

14 Buniello, A. et al. Open Targets Platform: facilitating therapeutic hypotheses building in drug discovery. Nucleic Acids Res 53, D1467–D1475 (2025). 10.1093/nar/gkae1128

15 Singh, T. et al. Rare coding variants in ten genes confer substantial risk for schizophrenia. Nature 604, 509–516 (2022). 10.1038/s41586-022-04556-w

16 Trubetskoy, V. et al. Mapping genomic loci implicates genes and synaptic biology in schizophrenia. Nature 604, 502–508 (2022). 10.1038/s41586-022-04434-5

17 Wickham, H. ggplot2: Elegant Graphics for Data Analysis. (Springer-Verlag, 2016).

18 Kramer, A., Green, J., Pollard, J., Jr. & Tugendreich, S. Causal analysis approaches in Ingenuity Pathway Analysis. Bioinformatics 30, 523–530 (2014). 10.1093/bioinformatics/btt703

19 Doncheva, N. T., Morris, J. H., Gorodkin, J. & Jensen, L. J. Cytoscape StringApp: Network Analysis and Visualization of Proteomics Data. Journal of Proteome Research 18, 623–632 (2019). 10.1021/acs.jproteome.8b00702

20 dplyr: A Grammar of Data Manipulation v. 1.1.4 (R package, 2023).

21 Johnson, J. L. et al. An atlas of substrate specificities for the human serine/threonine kinome. Nature 613, 759–766 (2023). 10.1038/s41586-022-05575-3

22 enrichplot: Visualization of Functional Enrichment Result v. 1.28.4 (Bioconductor, 2025).

23 Yu, G., Wang, L. G., Han, Y. & He, Q. Y. clusterProfiler: an R package for comparing biological themes among gene clusters. OMICS 16, 284–287 (2012). 10.1089/omi.2011.0118

24 Kick, F. L., Holze, H., Rydbirk, R. & Khodosevich, K. CRMetrics - an R package for Cell Ranger Filtering and Metrics Visualisation. (2023). 10.21203/rs.3.rs-2853524/v1

25 Wolock, S. L., Lopez, R. & Klein, A. M. Scrublet: Computational Identification of Cell Doublets in Single-Cell Transcriptomic Data. Cell Syst 8, 281–291 e289 (2019). 10.1016/j.cels.2018.11.005

26 Barkas, N. et al. Joint analysis of heterogeneous single-cell RNA-seq dataset collections. Nat Methods 16, 695–698 (2019). 10.1038/s41592-019-0466-z

27 pagoda2: Single Cell Analysis and Differential Expression v. 1.0.13 (R package, 2021).

28 Hao, Y. et al. Integrated analysis of multimodal single-cell data. Cell 184, 3573–3587 e3529 (2021). 10.1016/j.cell.2021.04.048

29 R: A Language and Environment for Statistical Computing (R Foundation for Statistical Computing, 2021).

30 gplots: Various R Programming Tools for Plotting Data v. 3.3.0 (R package, 2025).

31 Trapnell, C. et al. The dynamics and regulators of cell fate decisions are revealed by pseudotemporal ordering of single cells. Nat Biotechnol 32, 381–386 (2014). 10.1038/nbt.2859

