## Supplementary material for "Protein-state dysregulation and sex-specific neurodevelopmental signatures in schizophrenia forebrain organoids"

**Supplementary figures:**

**Figure S1:** Induced pluripotent stem cell (iPSC) line quality control (QC)

**Figure S2:** Dorsal forebrain organoid (FO) differentiation QC – CTL lines

**Figure S3:** FO differentiation QC – SCZ lines

**Figure S4:** Comparable growth and morphology of CTL and SCZ FOs during differentiation.

**Figure S5:** Single-nuclei RNA sequencing QC and analysis

**Figure S6:** Single-nuclei RNA sequencing of female and male samples separately

**Figure S7:** Analysis of postmortem cortical SCZ and CTL proteomics data from *Koopmans et. al 2026*

**Tables:**

**Table S1:** Patient-derived iPSC lines included in the study.

**Table S2:** Demographic distribution of iPSC cohort.

**Table S3:** Antibodies used for immunofluorescence staining.

**Table S4:** Confocal imaging acquisition details.

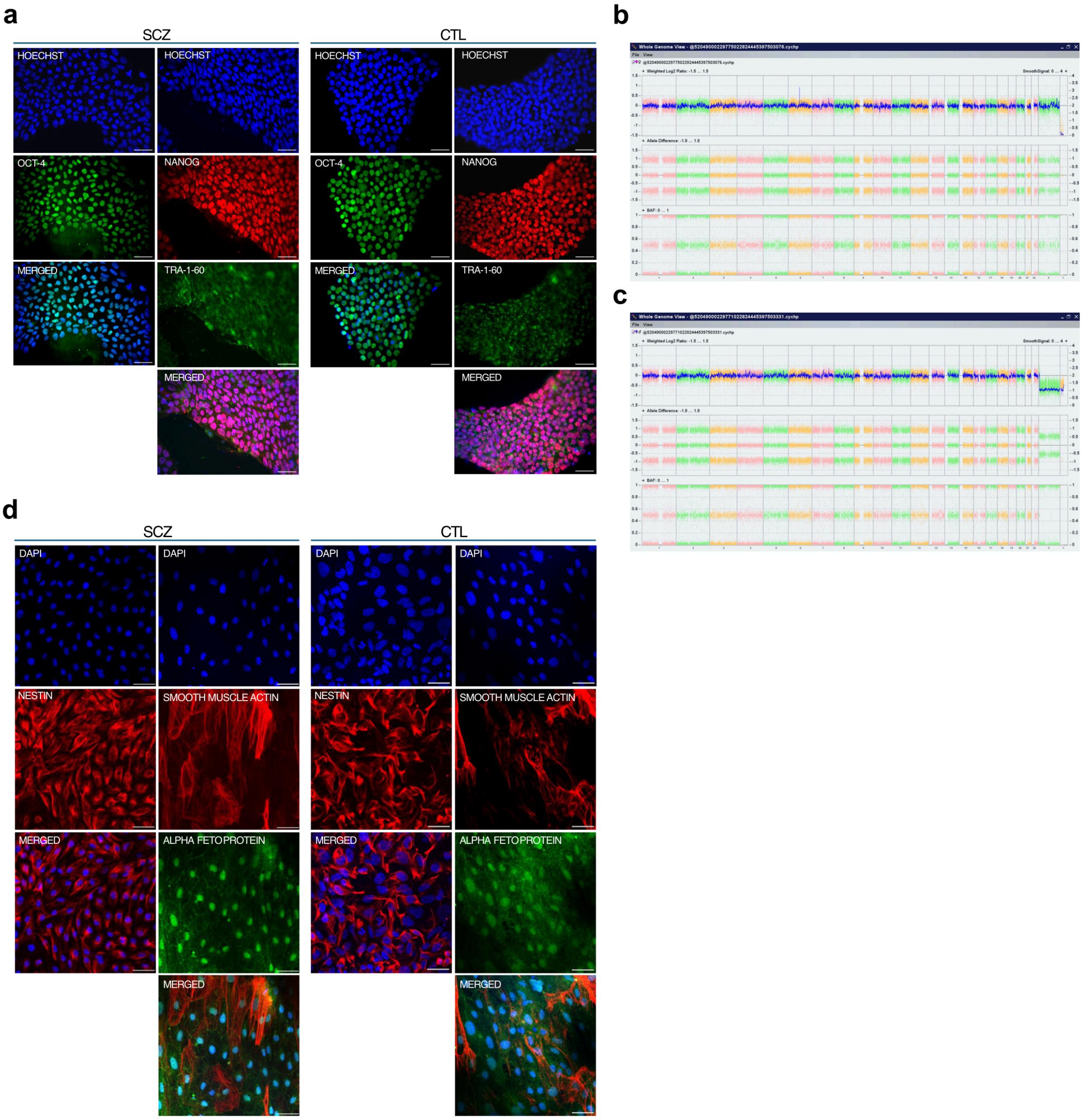


**Figure S1: Induced pluripotent stem cell (iPSC) line quality control (QC). a**. Representative confocal images of schizophrenia (SCZ) and control (CTL) iPSC lines showing expression of the pluripotency markers OCT4 (green), NANOG (red), and TRA-1-60 (green). Nuclei were stained with Hoechst (blue). **b-c.** Representative copy number variation (CNV) analysis of hiPSC lines (b; SCZ, c; CTL) using the CytoScan 750K array, demonstrating genomic integrity of the generated clones. **d.** Assessment of differentiation potential by embryoid body formation followed by spontaneous differentiation. Immunocytochemical staining confirmed expression of lineage-specific markers representing the three germ layers: nestin (ectoderm, red), smooth muscle actin (mesoderm, red), and alpha-fetoprotein (endoderm, green). Nuclei were counterstained with DAPI (blue).

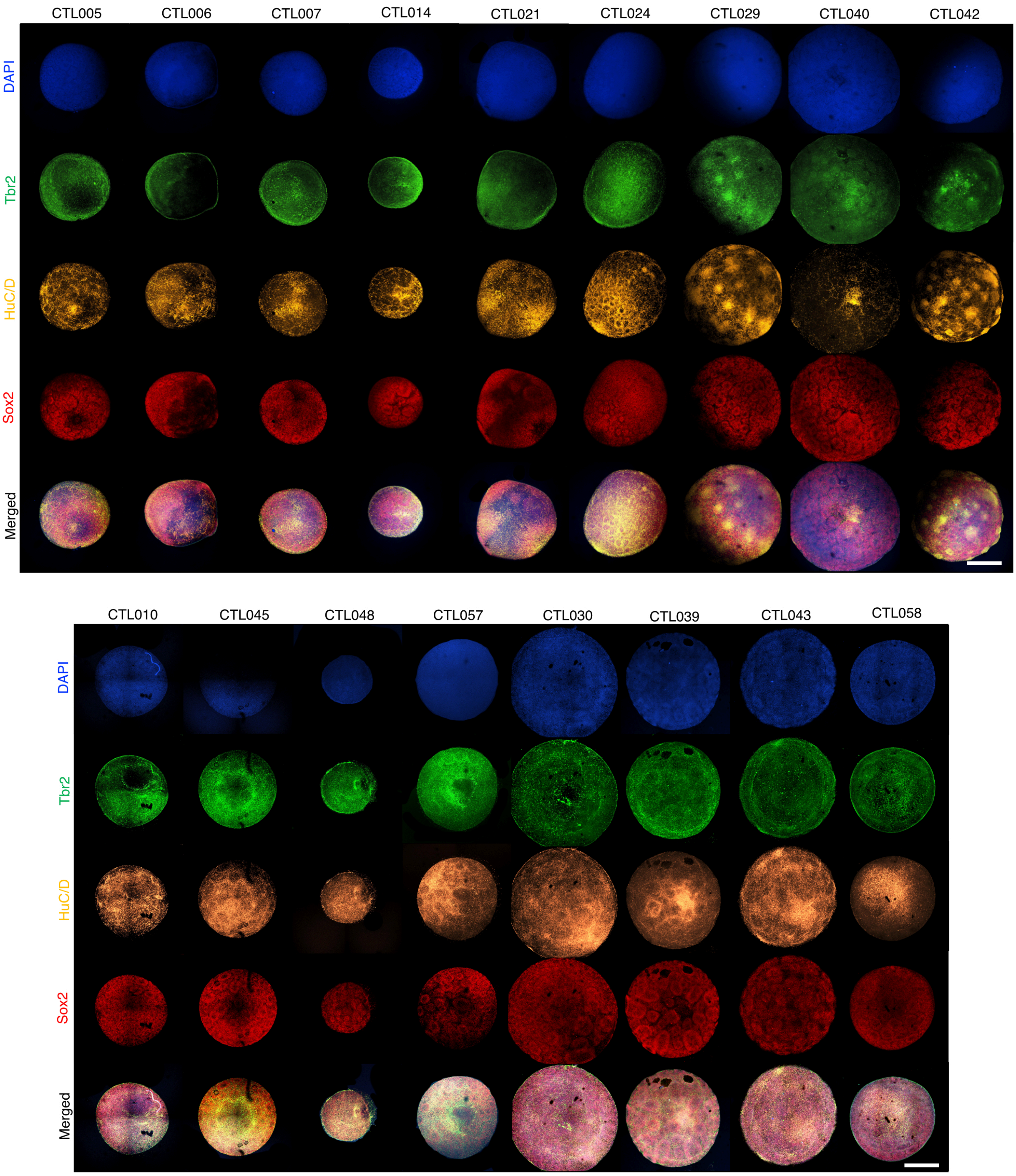


**Figure S2: Dorsal forebrain organoid (FO) differentiation QC – CTL lines.** Representative confocal images of all 17 CTL lines stained for TBR2 (green), HuC/D (yellow), SOX2 (red), and DAPI (blue) at day 20 of differentiation. Scalebar = 250 μm. All lines exhibited robust neuroepithelial organization and consistent dorsal forebrain patterning.

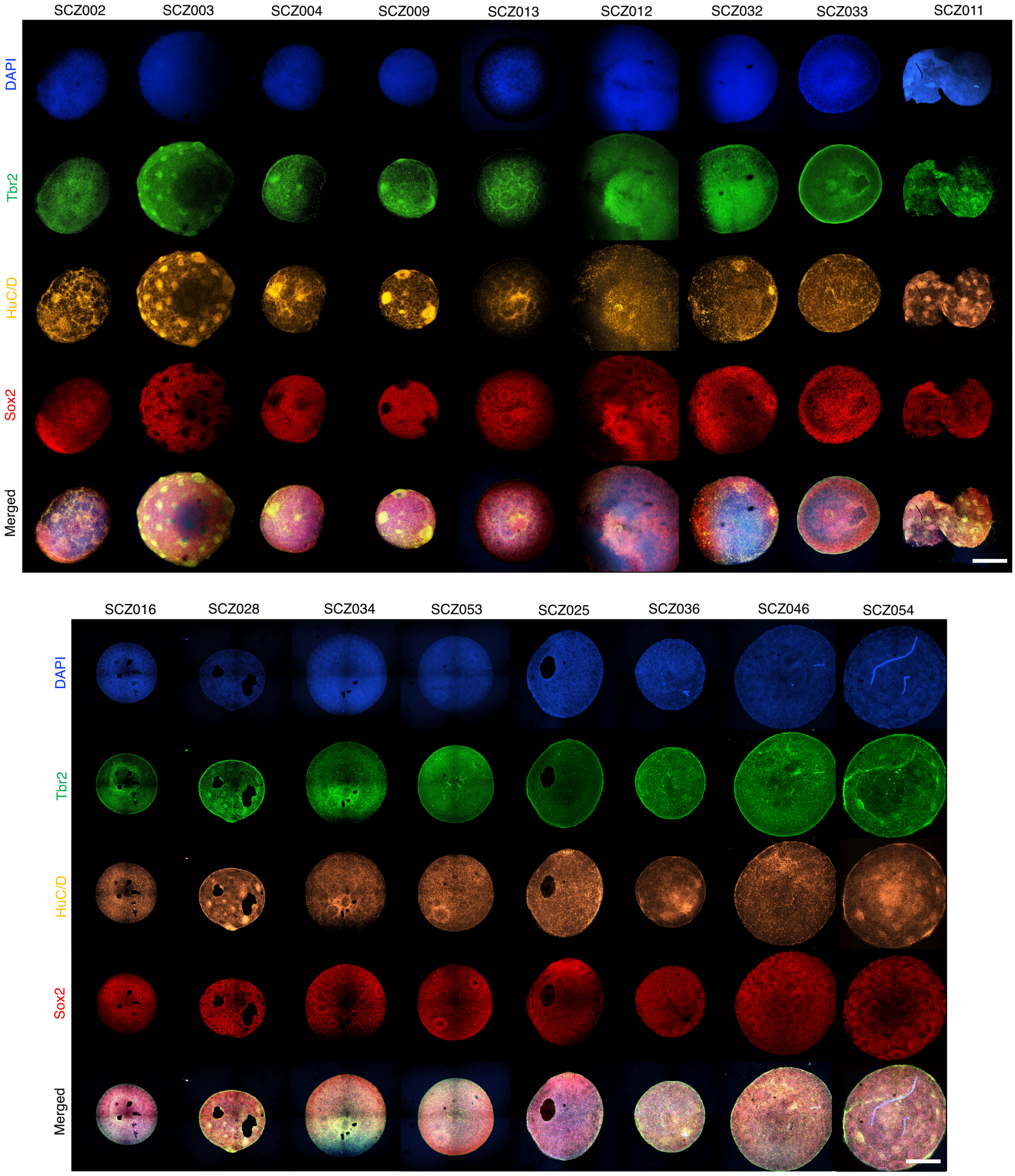


**Figure S3: FO differentiation QC – SCZ lines.** Representative confocal images of all 17 SCZ lines stained for TBR2 (green), HuC/D (yellow), SOX2 (red), and DAPI (blue) at day

20 of differentiation. Scalebar = 250 µm. All lines exhibited robust neuroepithelial organization and consistent dorsal forebrain patterning.

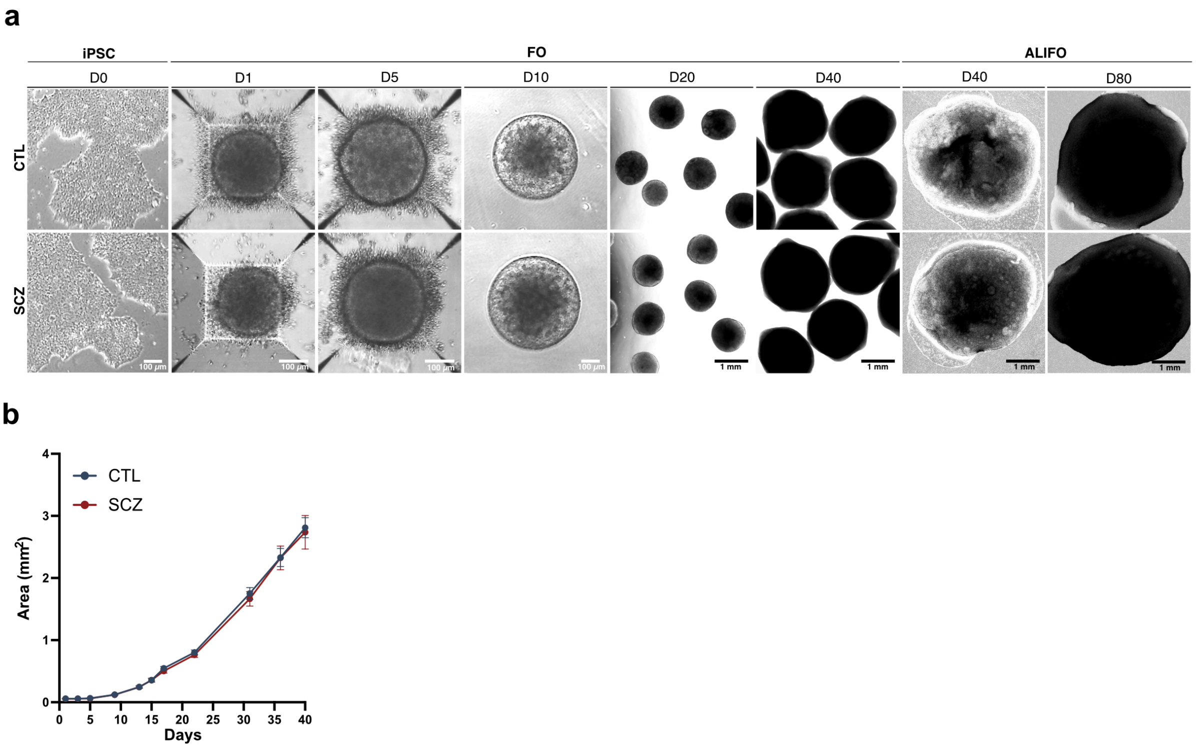


**Figure S4: Comparable growth and morphology of CTL and SCZ FOs during differentiation. a.** Representative brightfield images of CTL and SCZ cell lines across FO differentiation from induced pluripotent stem cell (iPSC) cultures (D0) through developing FOs to mature air-liquid interface FO cultures (ALIFOs) (D80). Scalebar = 100 µm (D0-D10) and 1 mm (D20-D80). **b.** Quantification of organoid area during differentiation from D0-40. n = 17 CTL and 17 SCZ, at least 6 FOs per replicate. Data are presented as mean ± SEM.

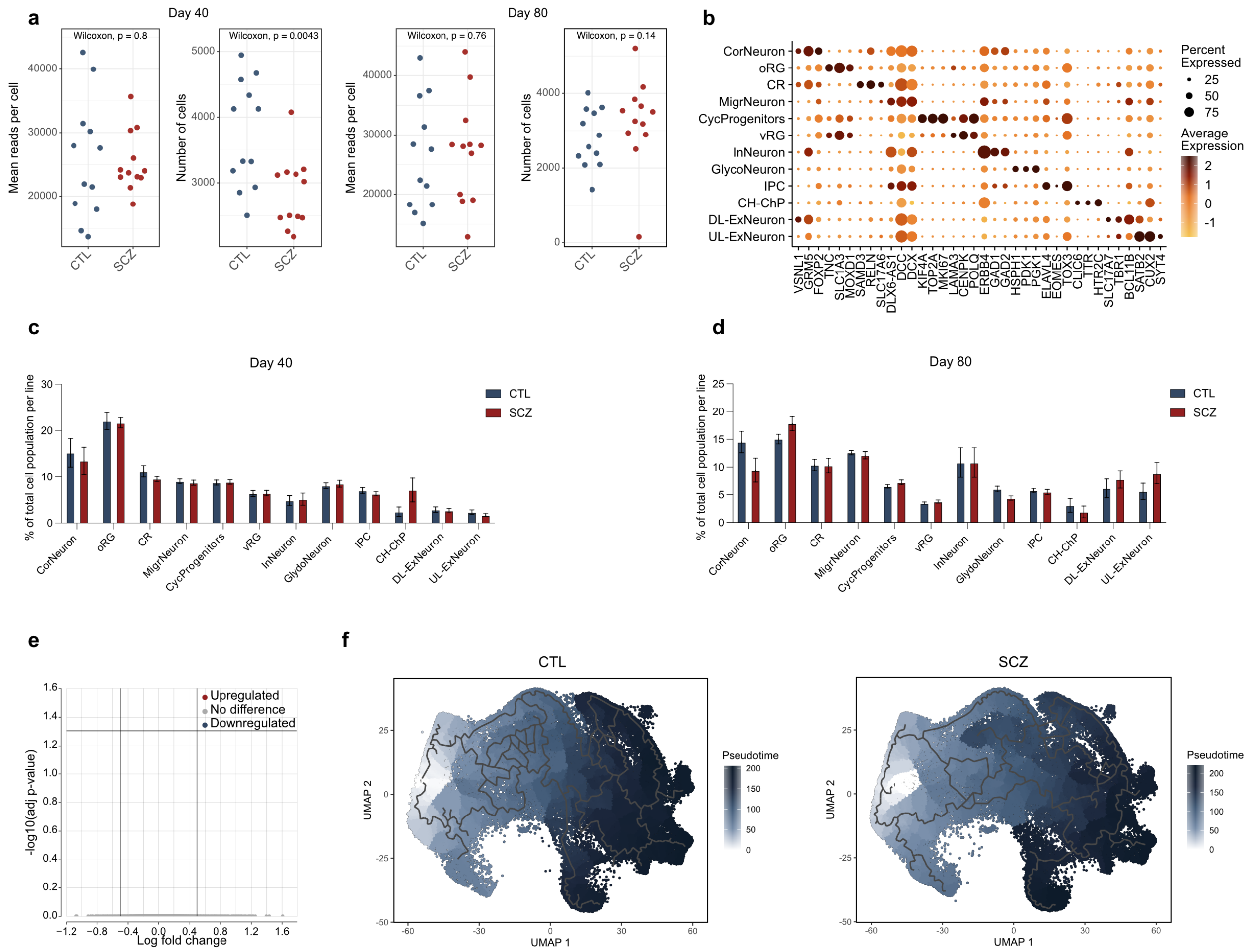

**Figure S5: Single-nuclei RNA sequencing (snRNAseq) QC and analysis. a.** mean reads per cell and number of cells per line at day 40 and day 80 from schizophrenia (SCZ) and control (CTL) forebrain organoids (FOs) and air-liquid interface forebrain organoids (ALIFOs). **b**. Expression of cell type-specific markers for each cluster with size of dot indicating the percentage of cluster cells expressing the marker and color of the dot the average expression log2 fold change compared to all cells not within the cluster. Cortical neurons (CorNeuron), outer radial glia (oRG), Cajal Retzius cells (CR), migratory neurons (MigrNeuron), cycling progneitors (CycProgenitors), ventricular radial glia (vRG), inhibitory neurons (InNeuron), glycolytic neurons (GlycoNeuron), intermediate progenitors (IPC), cortical hem/choroid plexus (CH-ChP), deep layer excitatory neurons (DL-ExNeuron), upper layer excitatory neurons (UL-ExNeuron). **c-d**. The percentage of cells within each cluster out of the total population for SCZ and CTL at c. day 40 and d. day 80. Data presented as mean ± SEM, two-tailed unpaired t-test. **e.** Volcanoplot of day 40 differentially expressed genes between SCZ/CTL. Limma voom-calculated adjusted p-value <0.05 and log-fold change >0.5 considered significant. **f**. Pseudotime analysis on CTL and SCZ, based on combined day 40 and -80 snRNAseq data, showing the developmental trajectories.

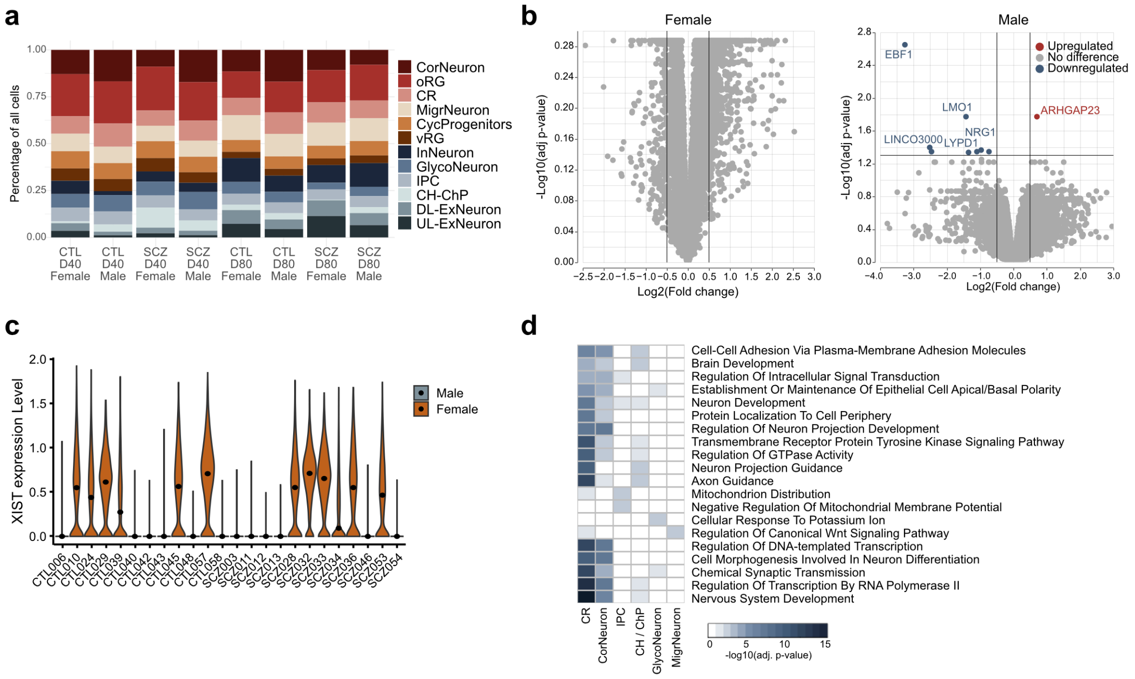

**Figure S6: Single-nuclei RNA sequencing of female and male samples separately. a.** Stacked barplot of day 40 and day 80 organoid single-nuclei RNA sequencing (snRNAseq) data showing the proportion of each cell population for schizophrenia (SCZ) and control (CTL) organoids per time point. Cortical neurons (CorNeuron), outer radial glia (oRG), Cajal Retzius cells (CR), migratory neurons (MigrNeuron), cycling progneitors (CycProgenitors), ventricular radial glia (vRG), inhibitory neurons (InNeuron), glycolytic neurons (GlycoNeuron), intermediate progenitors (IPC), cortical hem/choroid plexus (CH-ChP), deep layer excitatory neurons (DL-ExNeuron), upper layer excitatory neurons (UL-ExNeuron). **b.** Volcanoplots of day 80 differentially expressed genes (DEGs) between female and male SCZ/CTL. Limma voom-calculated adjusted p-value <0.05 and log-fold change >0.5 considered significant. **c.** Violin plots of XIST expression levels in all samples across the two timepoints (n = 12). **d.** Heatmap of top 20 enriched biological processes (BP) based on the six cell populations with significant DEGs in male SCZ/CTL. Fisher's exact test with Benjamini-Hochberg corrected p-value.

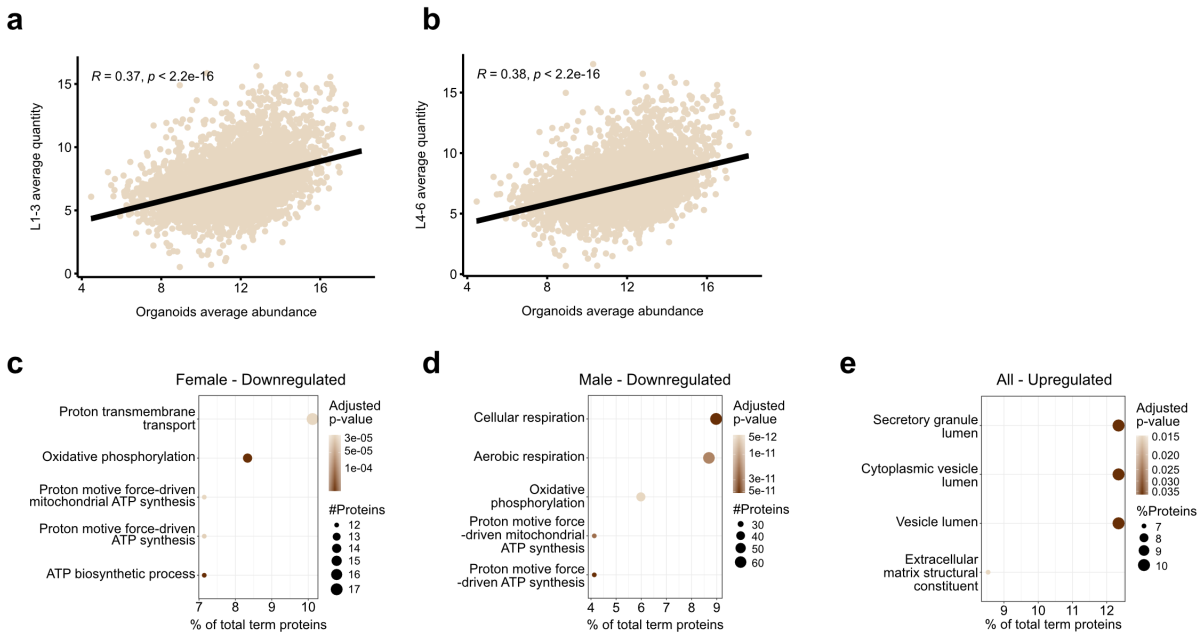


**Figure S7: Analysis of postmortem cortical SCZ and CTL proteomics data from *Koopmans et al. 2026*. a-b**. Correlation plots of the average protein abundance in forebrain organoids across all day 80 SCZ and CTL compared to the average protein quantity in a. layer 1-3 or b. layer 4-6 across all postmortem SCZ and CTL samples. Spearman's rank correlation. **c-d**. The top five GO terms enriched based on the significantly downregulated differentially expressed proteins (DEPs) in c. female and d. male and the significantly upregulated DEPs in all postmortem SCZ vs. CTL cortex with node colour indicating enrichment -log10(adjusted p-value), size the number of term-related DEPs and the % of total term proteins on the x-axis. Fisher's exact test with Benjamini-Hochberg correction.

| Cell line | Phenotype | Sex | Age |
|---|---|---|---|
| 005 | CTL | Female | 19 |
| 006 | CTL | Male | 28 |
| 007 | CTL | Male | 23 |
| 010 | CTL | Female | 20 |
| 014 | CTL | Female | 20 |
| 021 | CTL | Male | 22 |
| 024 | CTL | Female | 23 |
| 029 | CTL | Female | 26 |
| 030 | CTL | Male | 28 |
| 039 | CTL | Female | 23 |
| 040 | CTL | Male | 26 |
| 042 | CTL | Male | 28 |
| 043 | CTL | Male | 25 |
| 045 | CTL | Female | 25 |
| 048 | CTL | Male | 21 |
| 057 | CTL | Female | 25 |
| 058 | CTL | Male | 25 |
| 002 | SCZ | Male | 28 |
| 003 | SCZ | Male | 21 |
| 004 | SCZ | Female | 20 |
| 009 | SCZ | Female | 21 |
| 011 | SCZ | Male | 20 |
| 012 | SCZ | Male | 21 |
| 013 | SCZ | Male | 20 |
| 016 | SCZ | Male | 26 |
| 025 | SCZ | Male | 26 |
| 028 | SCZ | Female | 20 |
| 032 | SCZ | Female | 25 |
| 033 | SCZ | Female | 26 |
| 034 | SCZ | Female | 26 |
| 036 | SCZ | Female | 25 |
| 046 | SCZ | Male | 29 |
| 053 | SCZ | Female | 27 |
| 054 | SCZ | Male | 29 |

**Table S1: Patient-derived induced pluripotent stem cell (iPSC) lines included in the study.** Overview of control (CTL) and schizophrenia (SCZ) iPSC lines used for forebrain organoid generation, including donor phenotype, sex, and age at inclusion.

| Sex | Number | Age (mean ± SD) |
|---|---|---|
| Male | CTL: 9<br>SCZ: 9 | CTL: 25.11 ± 2.67<br>SCZ: 24.44 ± 3.91 |
| Female | CTL: 8<br>SCZ: 8 | CTL: 22.63 ± 2.67<br>SCZ: 23.75 ± 2.92 |

**Table S2: Demographic distribution of iPSC cohort.** Distribution of sex and age among control (CTL) and schizophrenia (SCZ) donor-derived iPSC lines included in the study. Male and female donors were equally represented across groups (CTL: n = 9 males, n = 8 females; SCZ: n = 9 males, n = 8 females). Donor age is presented as mean ± SD.

| Antibody | Company | Cat. No. | Dilution |
|---|---|---|---|
| Rabbit anti-OCT4 | Thermo Fischer Scientific | 701756 | 1:500 |
| Rabbit anti-SOX2 | Abcam | Ab97959 | 1:500 |
| Rabbit anti-homer1 | Synaptic systems | 160003 | 1:250 |
| Rabbit anti-FOXG1 | Thermo Fischer Scientific | 702554 | 1:250 |
| Rabbit anti-Doublecortin | Cell Signaling | 4604 | 1:500 |
| Rabbit anti-TBR1 | Abcam | Ab183032 | 1:200 |
| Rabbit anti-VGLUT1 | Synaptic systems | 135303 | 1:500 |
| Mouse anti-β-tubulin III | Sigma | T8660 | 1:2000 |
| Mouse anti-HuCD | Life Technologies | A21271 | 1:500 |
| Mouse anti-MAP2 | Sigma | M1406 | 1:2000 |
| Mouse anti-Nestin | BD Biosciences | BD611658 | 1:500 |
| Mouse anti-GAD67 | Merck Millipore | MAB5406 | 1:500 |
| Rat anti-CTIP2 | Abcam | Ab18465 | 1:500 |
| Guinea pig anti-synapsin1/2 | Synaptic systems | 106004 | 1:250 |
| Sheep anti-TBR2 | R&D Systems | AF6166 | 1:200 |
| Goat anti-Vimentin | R&D Systems | AF2105 | 1:200 |
| Chicken anti-β-tubulin III | Sigma | AB9354 | 1:1000 |
| Alexa Fluor 488 donkey anti-sheep | Invitrogen | A21202 | 1:1000 |
| Alexa Fluor 488 donkey anti-mouse | Invitrogen | A11015 | 1:500 |
| Alexa Fluor 488 goat anti-rabbit | Invitrogen | A11008 | 1:500 |
| Alexa Fluor 488 goat anti-mouse | Invitrogen | A11001 | 1:1000 |
| Alexa Fluor 488 goat anti-guinea pig | Abcam | Ab150185 | 1:500 |
| Alexa Fluor 568 donkey anti-goat | Invitrogen | A11057 | 1:1000 |
| Alexa Fluor 568 goat anti-mouse | Invitrogen | A11004 | 1:1000 |
| Alexa Fluor 568 goat anti-rat | Invitrogen | A11077 | 1:1000 |
| Alexa Fluor 647 donkey anti-rabbit | Invitrogen | A31573 | 1:1000 |
| Alexa Fluor 647 goat anti-rabbit | Invitrogen | A21245 | 1:1000 |

**Table S3: Antibodies used for immunofluorescence staining.** List of primary and secondary antibodies used for immunofluorescence staining, including host species, supplier information, catalog number (Cat. No.), and working dilution.

| Staining | Objective | Zoom | Resolution | Focal plane | z-stack details |
|---|---|---|---|---|---|
| OCT4/DAPI | X10 | - | 2560x2160 | Nuclei in focus | - |
| SOX2/TBR2/HuCD/DAPI | X10 | - | 2048x2048 | $SOX2^{+}$-rosettes in focus | - |
| Nestin/Vimentin/DCX/DAPI | X20 | X2 | 1024x1024 | Middle of rosettes (largest lumen) in focus | ± 5 µm above and below focus plane, step size 0.5 µm |
| SOX2/CTIP2/TBR1/DAPI | X10 | - | 2048x2048 | 50 µm from sample top | - |
| MAP2/CTIP2/FOXG1/DAPI | X10 | - | 2048x2048 | Staining in focus | 30 µm, step size 5 µm |
| GAD67/VGLUT1/BTUB/DAPI | X10 | X3 | 2048x2048 | Staining in focus | - |

**Table S4: Confocal imaging acquisition details.** Overview of confocal microscopy acquisition settings used for immunofluorescence imaging, including staining combination, objective, zoom factor, image resolution, focal plane selection, and z-stack acquisition parameters.
